# Casein Kinase II is Required for Proper Cell Division and Acts as a Negative Regulator of Centrosome Duplication in *C. elegans* Embryos

**DOI:** 10.1101/083378

**Authors:** Jeffrey C. Medley, Megan M. Kabara, Michael D. Stubenvoll, Lauren E. DeMeyer, Mi Hye Song

## Abstract

**Summary statement:** The conserved protein kinase CK2 negatively regulates centrosome assembly and is required for proper cell cycle progression and cytokinesis in early *C. elegans* embryos.

**Abstract:** Centrosomes are the primary microtubule-organizing centers that orchestrate microtubule dynamics during the cell cycle. The correct number of centrosomes is pivotal for establishing bipolar mitotic spindles that ensure accurate segregation of chromosomes. Thus, centrioles must duplicate once per cell cycle, one daughter per mother centriole, the process of which requires highly coordinated actions among core factors and modulators. Protein phosphorylation is shown to regulate the stability, localization and activity of centrosome proteins. Here, we report the function of Casein Kinase II (CK2) in early *C. elegans* embryos. The catalytic subunit (KIN-3/CK2α) of CK2 localizes to nuclei, centrosomes and midbodies. Inactivating CK2 leads to cell division defects, including chromosome missegregation, cytokinesis failure and aberrant centrosome behavior. Furthermore, depletion or inhibiting kinase activity of CK2 results in elevated ZYG-1 levels at centrosomes, restoring centrosome duplication and embryonic viability to *zyg-1* mutants. Our data suggest that CK2 functions in cell division and negatively regulates centrosome duplication in a kinase-dependent manner.

## Introduction

Control of proper centrosome number is crucial for the fidelity of cell division (Gonczy, 2015). In animal cells, centrosomes organize microtubules to direct the formation of bipolar mitotic spindles that contribute to accurate segregation of genomic content. Centrosomes comprise two orthogonally arranged centrioles surrounded by a dense network of proteins termed pericentriolar material (PCM). Centrioles must duplicate exactly once per cell cycle to provide daughter cells with the correct number of centrosomes. Cells with abnormal centrosome number are prone to errors in DNA segregation and cytokinesis, leading to genomic instability and tumorigenesis (Godinho and Pellman, 2014).

Genetic analyses in *C. elegans* have elucidated a core set of five conserved factors required for centriole duplication, including the Plk4 related kinase ZYG-1 (O’Connell et al., 2001), the coiled coil protein SPD-2 (Kemp et al., 2004, Pelletier et al., 2004), SAS-5 (Delattre et al., 2004), SAS-6 (Leidel et al., 2005) and SAS-4 (Kirkham et al., 2003, Leidel and Gonczy, 2003). SPD-2 and ZYG-1 localize early to the site of nascent centriole formation and are required for the sequential recruitment of the SAS-5/SAS-6 complex and SAS-4 (Delattre et al., 2006, Pelletier et al., 2006). In human cells, increased activity of centriole factors (Plk4, STIL or HsSAS6) leads to centrosome amplification (Arquint and Nigg, 2014, Kleylein-Sohn et al., 2007, Strnad et al., 2007), suggesting that regulating the levels and activity of those factors is essential for precise control of centrosome number.

Post-translational modifications have been also implicated in centrosome assembly. For example, protein phosphorylations regulate the localization and stability of core centriole regulators. In humans and flies, the kinase Plk4 is required for recruiting STIL/Ana2/SAS-5 and SAS-6 to nascent centrioles via phosphorylation (Dzhindzhev et al., 2014, Kratz et al., 2015). Plk4 levels are controlled through autophosphorylation followed by proteosomal degradation (Cunha-Ferreira et al., 2013, Guderian et al., 2010, Holland et al., 2010, Klebba et al., 2013). Many other protein kinases are also shown to control centrosome assembly and function, including PLK-1 (Decker et al., 2011, Lane and Nigg, 1996), Aurora A Kinase (Hannak et al., 2001), and CDK2/Cyclin E (Hinchcliffe et al., 1999).

Conversely, Protein Phosphatase 2A (PP2A) plays a key role in centrosome duplication, and such role appears to be conserved in humans, flies and nematodes (Brownlee et al., 2011, Kitagawa et al., 2011, Song et al., 2011), underlying the coordinated kinase/phosphatase action in regulating centrosome duplication. The *C. elegans* catalytic subunit of PP2A, LET-92, was identified by proteomic analysis of the RNA-binding protein SZY-20 that negatively regulates a core centriole factor ZYG-1 (Song et al., 2008, Song et al., 2011). It has been shown that PP2A positively regulates centrosome duplication by promoting the stability and/or localization of ZYG-1 and SAS-5 in the *C. elegans* embryo (Kitagawa et al., 2011, Song et al., 2011). However, the counteracting kinase(s) to PP2A in centrosome duplication has not yet been identified. Interestingly, the proteomic approach that identified SZY-20 associating factors has listed both positive and negative regulators of centrosome duplication (Song et al., 2011, Stubenvoll et al., 2016). Thus, it is possible that one or more kinases interacting with SZY-20 might counteract PP2A in centrosome assembly.

One of the SZY-20 interacting factors we identified is the *C. elegans* Casein Kinase 2 holoenzyme (CK2), an evolutionarily conserved serine/threonine protein kinase (Hu and Rubin, 1990). In general, CK2 acts as a tetrameric holoenzyme comprising two catalytic (CK2α) and two regulatory (CK2β) subunits (Niefind et al., 2009). CK2 targets a large number of substrates that are involved in cellular proliferation (Meggio and Pinna, 2003). Aberrant CK2α activity has been shown to result in centrosome amplification in mammalian cells (St-Denis et al., 2009). Consistently, elevated CK2 activities are frequently observed in many human cancers (Guerra and Issinger, 2008, Trembley et al., 2010). The *C. elegans* genome contains a single catalytic and regulatory subunit encoded by *kin-3* and *kin-10,* respectively (Hu and Rubin, 1990, Hu and Rubin, 1991). In *C. elegans*, CK2 is also shown to function in a broad range of cellular processes, including cell proliferation in the germ line (Wang et al., 2014), Wnt signaling (de Groot et al., 2014), male sensory cilia (Hu et al., 2006) and microRNA targeting (Alessi et al., 2015). However, while genome-wide RNAi screening indicated that CK2 is required for the viability of *C. elegans* embryos (Fraser et al., 2000, Sonnichsen et al., 2005), the function of CK2 during embryonic development remains largely undefined. In this study, we investigated the role of *C. elegans* CK2 during cell division in early embryos. We show that protein kinase CK2 is required for proper cell division and cytokinesis and negatively regulates centriole duplication.

## Results

We have identified KIN-3/CK2α, the catalytic subunit of protein kinase CK2 through mass spectrometry of co-immunoprecipitated materials with anti-SZY-20. The conserved RNA-binding protein SZY-20 is shown to negatively regulate centrosome assembly by opposing ZYG-1 at the centrosome (Kemp et al., 2007, Song et al., 2008). Mass spectrometry identified three KIN-3 peptides (11% coverage; **Table S1**) that were present in SZY-20 co-precipitates but absent in IgG control. Consistently, we also show that KIN-3::GFP co-precipitates with SZY-20 (10.4% coverage) and the regulatory subunit of CK2, KIN-10/CK2β (5.1% coverage) (**Table S1**). Further, we confirmed the physical interaction between KIN-3 and SZY-20 using immunoprecipitation followed by immunoblot (**Fig. S1**). Together, our results suggest that the protein kinase CK2 holoenzyme physically associates with SZY-20 *in vivo*.

### CK2 acts as a negative regulator of centrosome duplication

Given that *szy-20* is a known genetic suppressor of *zyg-1* (Kemp et al., 2007, Song et al., 2008), we asked if CK2 functions in centrosome assembly using *zyg-1(it25)* mutants*. zyg-1(it25)* is a temperature-sensitive (ts) mutation that, at the restrictive temperature 24°C, results in a failure of centrosome duplication, leading to monopolar spindles at the second mitosis and 100% embryonic lethality (O’Connell et al., 2001). At the semi-restrictive temperature 22.5°C, *kin-3(RNAi)* significantly increased (25 ± 25%) embryonic viability to *zyg-1(it25)* mutants compared to control RNAi (8.0 ±13%, *p* <0.001) (**Fig. 1A, Table S2),** while *kin-3(RNAi)* produced embryonic lethality (47 ± 10%, n=2433) to wild-type animals (**Table S2)**. In contrast, at the restrictive temperature 24°C, *kin-3(RNAi)* had no significant effect on embryonic viability of *zyg-1(it25)* mutants (**Fig. 1A)** although *kin-3(RNAi)* had a similar effect on embryonic lethality of wild-type animals (44 ± 16%, n=3724) (**Table S2)**. Thus, knocking down KIN-3 partially restores embryonic viability in *zyg-1(it25)* mutants, suggesting that *kin-3* acts as a genetic suppressor of *zyg-* 1, perhaps through ZYG-1 activity to regulate centrosome assembly.

**Figure 1.**
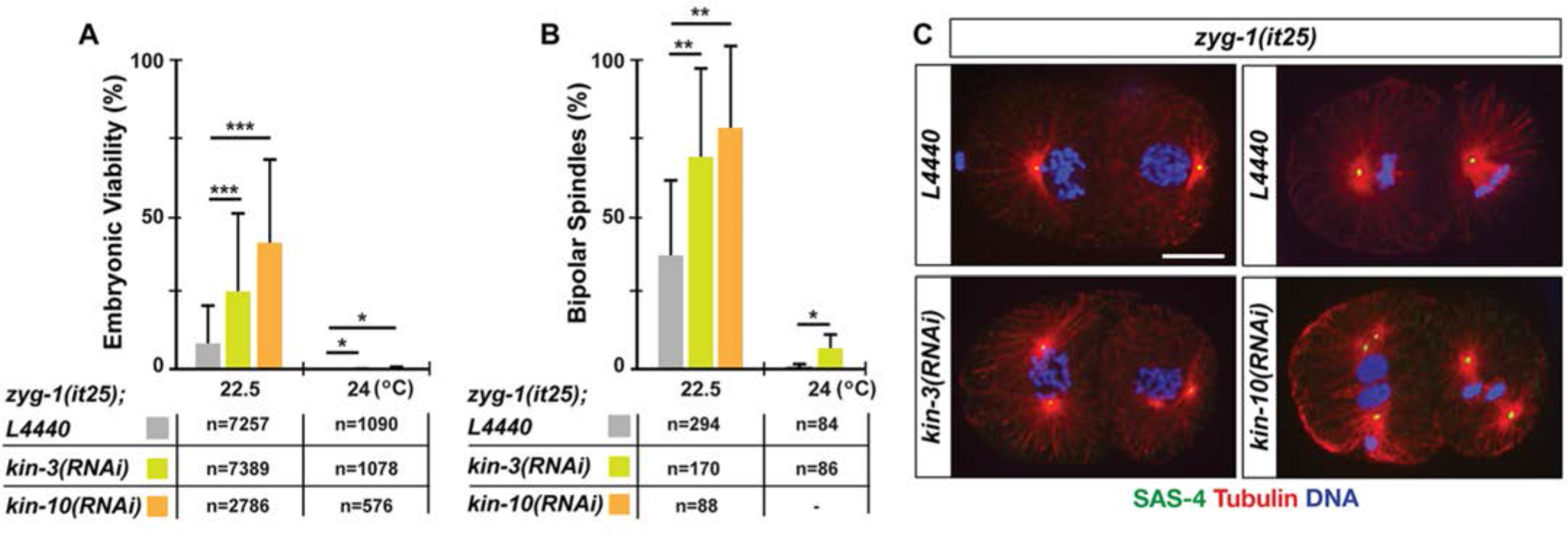
Knocking down CK2 partially restores embryonic viability and bipolar spindle formation to *zyg-1(it25)*. Depleting CK2 subunits by either *kin-3(RNAi)* or *kin-10(RNAi)* leads to an increase in both (A) embryonic viability and (B) bipolar spindle formation to *zyg-1(it25)* mutants, which were examined at restrictive (24°C) and semi-restrictive temperatures (21.5 and 22.5°C). (A,B) Average values are presented. Error bars are standard deviation (s.d). n is given as the number of embryos (A) or the number of centrosome duplication events (B) scored. \*\*\**p* <0.001, \*\**p* <0.01, \**p* <0.05 (two-tailed t-test). (C) Immunofluorescence of *zyg-1(it25)* embryos raised at 22.5°C illustrates mitotic spindles at second mitosis. *kin-3(RNAi)* or *kin-10(RNAi)* restores bipolar spindles to *zyg-1(it25)* embryos, but control embryos display monopoles. SAS-4 was used as a centriole marker. Scale bar, 10 μm.

As *zyg-1* is essential for centrosome duplication, it seems likely that *kin-3(RNAi)* restores embryonic viability to *zyg-1(it25)* by rescuing centrosome duplication. To test this, we examined *zyg-1(it25); kin-3(RNAi)* embryos that formed mitotic spindles at the second mitosis to quantify the event of successful centriole duplication during the first cell cycle. Compared to control RNAi, *kin-3(RNAi)* led to a significant fold increase in bipolar spindle formation to *zyg-1(it25)* embryos at all temperatures examined (**Fig. 1B,C**). Therefore, *kin-3(RNAi)* restores embryonic viability to *zyg-1(it25)* through restoration in centriole duplication. We then questioned if KIN-3 regulates centrosome duplication as part of the CK2 holoenzyme or as a free subunit, independently of the holoenzyme. If the holoenzyme CK2 functions in centrosome regulation, we should observe similar effects on *zyg-1(it25)* mutants by depleting KIN-10, the sole regulatory subunit (CK2β) of CK2 in *C. elegans* (**Fig. 1B,C**). Compared to control, *kin-10(RNAi)* resulted in significantly increased levels of embryonic viability and bipolar spindle formation to *zyg-1(it25)* embryos at the semi-restrictive temperature, 22.5°C. Thus, it is likely that the protein kinase CK2 holoenzyme has a role in centrosome duplication as a negative regulator.

Since KIN-3 physically interacts with SZY-20, we asked whether *kin-3* and *szy-20* genetically interact as well. We tested how co-depletion of KIN-3 and SZY-20 affects bipolar spindle formation in *zyg-1(it25)* embryos (**Fig. S2**). At the restrictive temperature, 24°C, the *szy-20(bs52)* mutation restores bipolar spindle formation to *zyg-1(it25)* embryos (49 ± 12%) as reported previously (Song et al., 2008). Strikingly, depleting KIN-3 in *zyg-1(it25);szy-20(bs52)* double mutants led to nearly 100% bipolar spindle formation to *zyg-1(it25)*, whereas *kin-3(RNAi)* alone exhibits much lower level (13 ± 8 %) of bipolar spindle formation in *zyg-1(it25).* Thus, co-depleting *kin-3* and *szy-20* enhances bipolar spindle formation to *zyg-1(it25)* mutants, which suggests that *kin-3* genetically interacts with *szy-20* to negatively regulate centrosome assembly.

### CK2 negatively regulates centrosomal ZYG-1 levels

How does protein kinase CK2 holoenzyme influence centrosome duplication? Given that protein phosphorylations are known to regulate the stability, localization and activity of centrosome factors (Brownlee et al., 2011, Cunha-Ferreira et al., 2013, Guderian et al., 2010, Holland et al., 2010, Klebba et al., 2013, Song et al., 2011), we first examined whether KIN-3 affected ZYG-1 localization to centrosomes in wild-type embryos. We immunostained embryos using anti-ZYG-1 (Stubenvoll et al., 2016), and quantified centrosomal levels of ZYG-1 in *kin-3(RNAi)* embryos compared to controls. We measured the fluorescence intensity of centrosome-associated ZYG-1 at the first anaphase where ZYG-1 levels at centrosomes are highest (Song et al., 2008). In embryos depleted of KIN-3, we observed a significant increase in centrosomal ZYG-1 levels (1.7 ± 0.7 fold; *p* <0.0001) compared to controls (**Fig. 2A,B**). We continued to examine the effect of KIN-3 knockdown on other centriole factors. By immunostaining embryos for SAS-4 (Kirkham et al., 2003, Leidel and Gonczy, 2003) and SAS-5 (Delattre et al., 2004), we quantified the levels of centrosome-associated SAS-5 and SAS-4 at first anaphase (**Fig. 2A,B**). However, we observed no significant changes in centrosomal levels of SAS-4 or SAS-5 between *kin-3(RNAi)* and control embryos.

**Figure 2.**
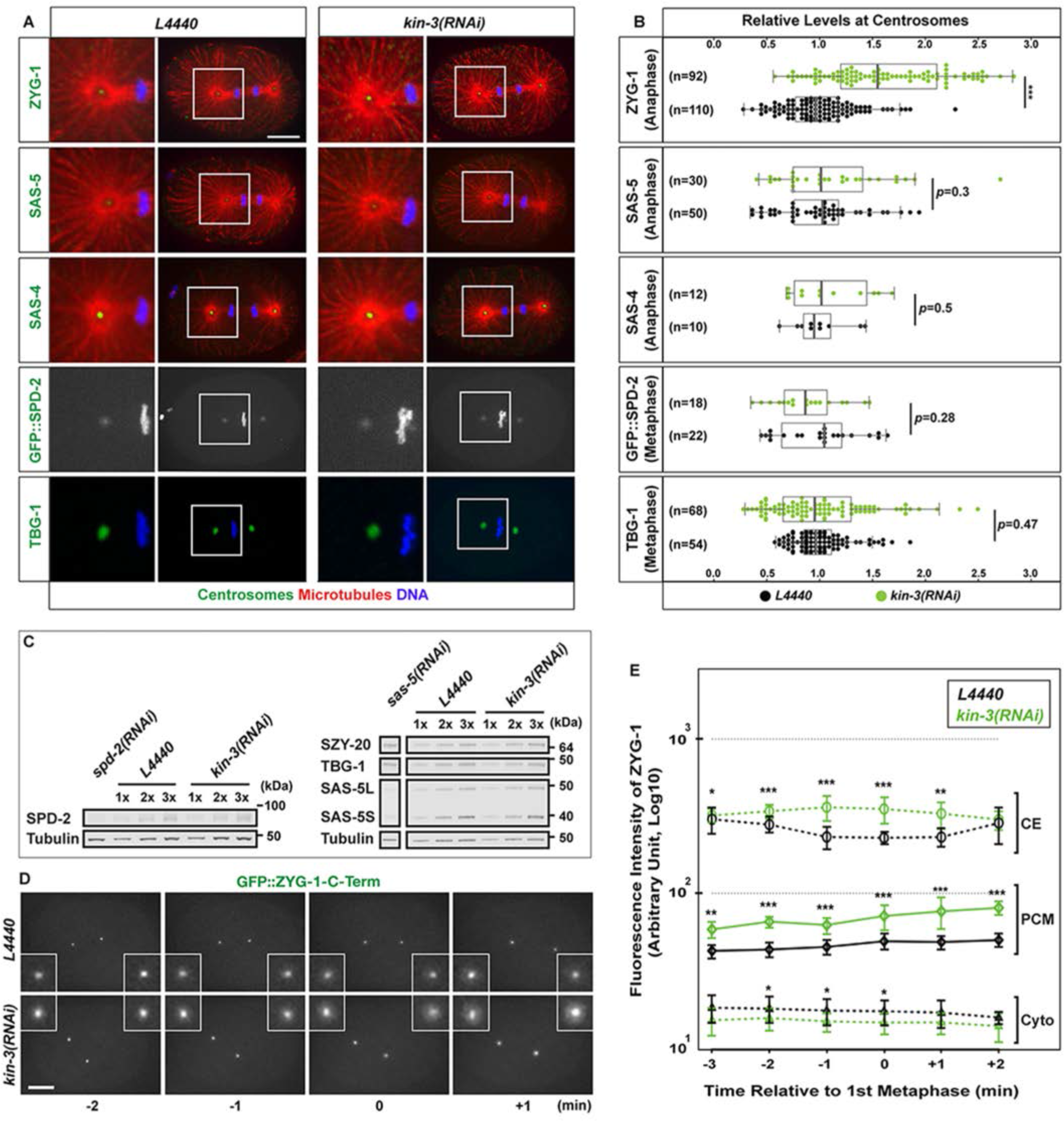
KIN-3 depletion leads to increased levels of centrosomal ZYG-1. (A) Wild-type embryos stained for centrosome factors (left) illustrate centrosomal localization of each factor. To assess the centriolar localization of ZYG-1, SAS-5 and SAS-4, embryos were co-stained for Tubulin to mark centrosomes. For SPD-2, shown are still-images from time-lapse movies of embryos expressing GFP::SPD-2 and GFP::Histone. (B) Quantification of centrosomal levels of each factor in *kin-3(RNAi)* (green dots) relative to controls (black dots). Each dot represents a centrosome. Box ranges from the first through third quartile of the data. Thick bar indicates the median. Dashed line extends 1.5 times the inter-quartile range or to the minimum and maximum data point. \*\*\**p* <0.001 (two-tailed t-test). (C) Quantitative immunoblot analyses of embryonic lysates reveal that KIN-3 knockdown leads to no significant changes in overall levels of SPD-2 (0.92 ± 0.41 fold, n=4), SZY-20 (1.03 ± 0.3, n=7), TBG-1 (1.06 ± 0.35, n=10), both isoforms of SAS-5L (404aa: 1.01 ± 0.21, n=16) and SAS-5S (288aa: 1.04 ± 0.42, n=15) relative to controls. n indicates the number of biological replicates. Tubulin was used as a loading control. (D) Still images from time-lapse movies of embryos expressing GFP::ZYG-1-C-term. Depleting KIN-3 results in elevated levels of centrosomal GFP::ZYG-1 throughout the first cell cycle. Time (min) is given relative to first metaphase (t=0). (E) Quantification of GFP::ZYG-1-C-term levels in *kin-3(RNAi)* and control embryos. Average fluorescence intensity is plotted (n=10 centrosomes in 5 embryos). Error bars are s.d. \*\*\**p* <0.001, \*\**p* <0.01, \**p* <0.05 (two-tailed t-test). (A,D) Insets highlight centrosomes magnified 4 folds. Scale bar, 10 μm.

SPD-2 localizes to both PCM and centrioles (Kemp et al., 2004, Pelletier et al., 2004) and is required for localization of ZYG-1 (Delattre et al., 2006, Pelletier et al., 2006). Increased levels of ZYG-1 at centrosomes could result from elevated SPD-2. To address this, we acquired time-lapse movies of KIN-3 depleted embryos expressing GFP::SPD-2 and GFP::histone (Kemp et al., 2004). Quantifying the fluorescence intensity of GFP::SPD-2 at first metaphase centrosomes showed no significant changes (0.88 ± 0.32 fold, *p* =0.3) in centrosomal SPD-2 levels in *kin-3(RNAi)* embryos compared to control (**Fig. 2A,B**). Thus, elevated ZYG-1 levels in *kin-3(RNAi)* centrosomes are unlikely due to SPD-2 activity. We also examined another PCM factor TBG-1 (γ-tubulin; Hannak et al., 2001). By quantitative immunofluorescence using anti-TBG-1, we observed no significant change in centrosomal TBG-1 levels between *kin-3(RNAi)* and control embryos. Further, quantitative immunoblot analyses revealed that overall levels of centrosome factors tested were unaffected by KIN-3 depletion (**Fig. 2C**).

Together, our results suggest that CK2 specifically regulates ZYG-1 levels at the centrosome. We then asked if increased ZYG-1 levels at first anaphase centrosomes resulted from a cell cycle shift by loss of KIN-3. To address cell cycle dependence, we recorded 4D time-lapse movies using embryos expressing GFP::ZYG-1-C-term that contains a C-terminal fragment (217-706 aa) of ZYG-1 and localizes to centrosomes (Peters et al., 2010, Shimanovskaya et al., 2014), starting from pronuclear migration through visual separation of centriole pairs at anaphase during the first cell cycle (**Fig. 2D, Movie S1).** Throughout the first cell cycle in *kin-3(RNAi)* embryos, we observed a 1.5 fold increase in total centrosome-associated (PCM-like) ZYG-1 levels relative to control (**Fig. 2E)**, but no significant difference in cytoplasmic levels between *kin-3(RNAi)* and control embryos (**Fig. 2E**), suggesting that KIN-3 influences centrosome-associated ZYG-1 levels throughout the cell cycle. However, we cannot exclude the possibility that KIN-3 might regulate overall ZYG-1 levels, and thereby influencing ZYG-1 levels at centrosomes. While direct measurement of endogenous levels of ZYG-1 will address this question, we were unable to assess overall levels of ZYG-1 due to technical limits to the detection of endogenous levels of ZYG-1, largely owing to low abundance of ZYG-1 in *C. elegans* embryos.

### Depletion of CK2 restores ZYG-1 levels at *zyg-1* mutant centrosomes

Elevated levels of centrosomal ZYG-1 by depleting CK2 might lead to the restoration of bipolar spindles and embryonic viability to *zyg-1* mutants. It has been shown that several genetic suppressors of *zyg-1* regulate ZYG-1 levels at centrosomes (Peel et al., 2012, Song et al., 2011, Stubenvoll et al., 2016). We show that centrosomes in *zyg-1(it25)* embryos exhibit a significantly reduced level (~60%) of ZYG-1 compared to wild-type controls at first anaphase (**Fig. 3A,B**). In fact, depleting CK2 by *kin-3(RNAi)* or *kin-10(RNAi)* resulted in significantly increased ZYG-1 levels at *zyg-1(it25)* centrosomes (**Fig. 3A,B**). Consistently, *kin-10(RNAi)* led to increased centrosomal ZYG-1 in wild-type embryos (**Fig. S3**), comparable to the effect by *kin-3(RNAi)* (**Fig. 2A,B**). Together, our results suggest that inhibiting the holoenzyme CK2 restores centrosome duplication to *zyg-1(it25),* at least in part, by increasing centrosome-associated ZYG-1 levels at centrosomes.

**Figure 3.**
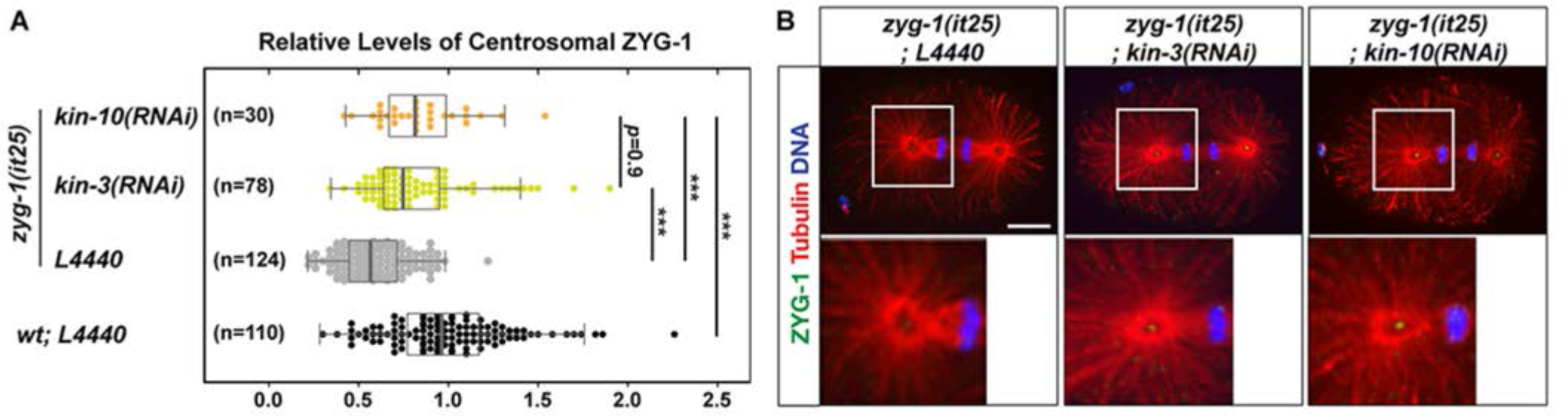
Depleting CK2 restores centrosomal ZYG-1 levels in *zyg-1(it25)* embryos. (A) Quantification of centrosomal ZYG-1 levels at first anaphase in wild-type and *zyg-1(it25)* embryos exposed to *kin-3(RNAi), kin-10(RNAi)* or *L4440*. Values are relative to wild-type centrosomes treated with control RNAi (*L4440*). At the semi-restrictive temperature 22.5°C, *zyg-1(it25)* embryos exhibit reduced ZYG-1 levels at centrosomes (0.59 ± 0.32) compared to the wild-type. *kin-3(RNAi)* or *kin-10(RNAi)* in *zyg-1(it25)* mutants restores centrosomal ZYG-1 levels to near wild-type levels (0.83 ± 0.50 and 0.84 ± 0.42, respectively). Each dot represents a centrosome. Box ranges from the first through third quartile of the data, and thick bar represents the median. Dashed line extends 1.5 times the inter-quartile range or to the minimum and maximum data point. \*\*\**p* <0.001 (two-tailed t-test). Note that data for control RNAi treated wild-type embryos are also presented in Fig 2B, but included for quantitative analysis. (B) *zyg-1(it25)* embryos stained for ZYG-1 illustrate centrosome-associated ZYG-1 localization. Inset illustrates centrosomal regions magnified 4 folds. Scale bar, 10 μm.

### CK2 is required for proper cell division in *C. elegans* embryos

Prior studies in *C. elegans* have shown that CK2 is required for embryonic viability (Fraser et al., 2000, Sonnichsen et al., 2005) and that CK2 functions in stem cell proliferation in germ line development (Wang et al., 2014). However, the specific role of CK2 in *C. elegans* embryos has not been examined in detail. Given both subunits of CK2 physically associate with SZY-20, we were interested to see what role CK2 plays in the early cell cycle. As homozygous mutant animals for *kin-3* or *kin-10* arrest at late larval stages, we treated L4 stage worms by RNAi-feeding to knockdown the catalytic subunit (KIN-3/CK2α) of CK2, and examined the knockdown effect in early embryos. As reported previously (Alessi et al., 2015), animals exposed to *kin-3(RNAi)* for 24 hours produced no significant embryonic lethality although these animals exhibited other phenotypes, such as sterility and protruding vulva (wormbase.org; Wang et al., 2014). Strikingly, when L4 worms were exposed for an extended period (36-48 hours) to *kin-3(RNAi)*, these animals produced significant and reproducible embryonic lethality (**Fig. 4A, Table S2**) and a reduction in brood-size (**Fig. 4B**). We also observed that *kin-10(RNAi)* led to a similar level of embryonic lethality (**Fig. 4A, Table S2**), suggesting that protein kinase holoenzyme CK2 is required for *C. elegans* embryogenesis.

**Figure 4.**
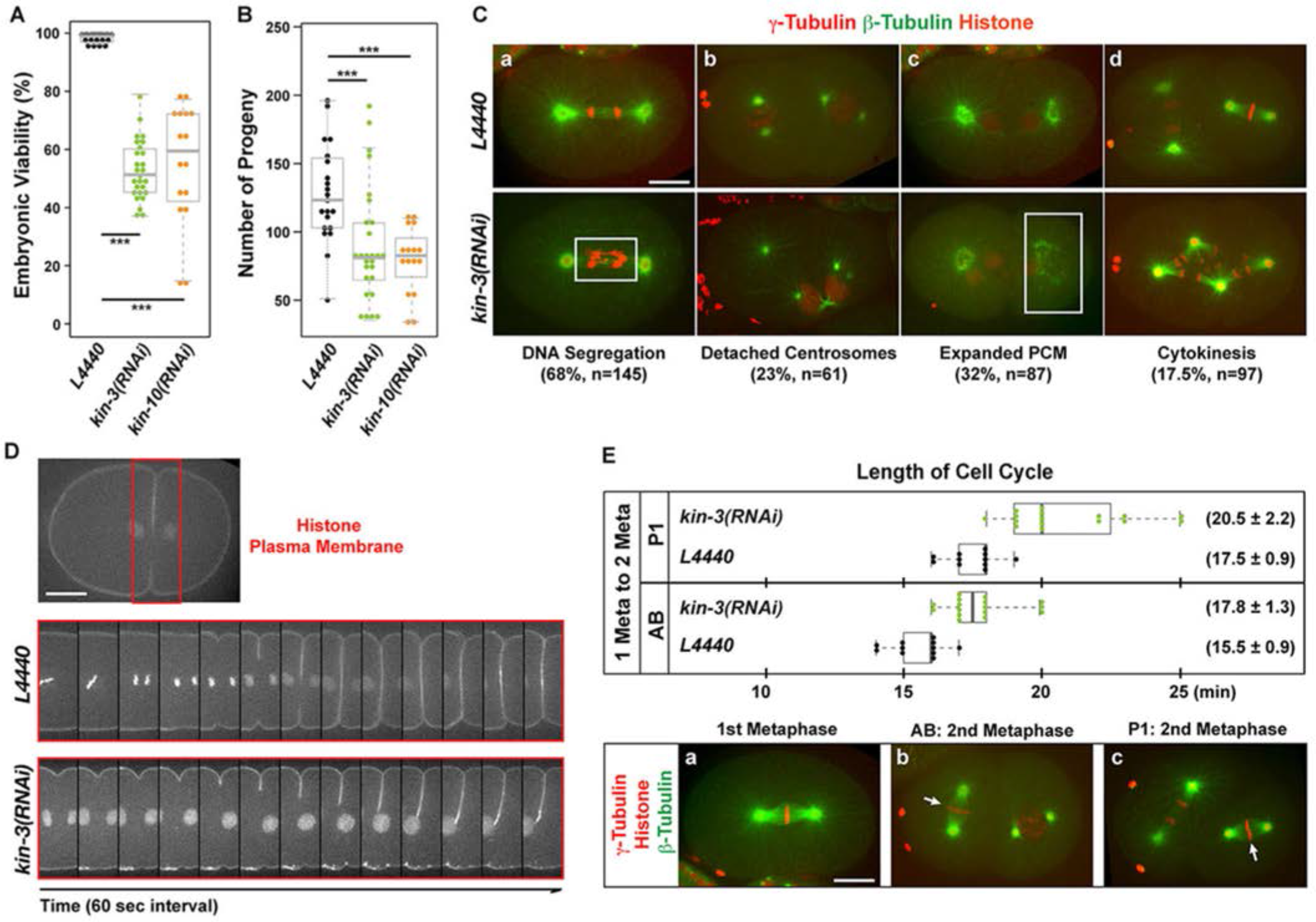
CK2 is required for early cell divisions in *C. elegans* embryos. (A) Knockdown of KIN-3 or KIN-10 by RNAi results in embryonic lethality (average ± s.d.: 47 ± 10% and 45 ± 21%, respectively). (B) Knockdown of KIN-3 or KIN-10 by RNAi leads to a significant reduction in the number of progeny (90 ± 43 and 79 ± 24, respectively) compared to controls (133 ± 43). (A,B) Each dot represents an animal. (C) Wild-type embryos expressing GFP::β-tubulin, mCherry::γ-tubulin and mCherry:histone: *kin-3(RNAi)* results in defective cell divisions including errors in (a) lagging DNA (box, **Movie S2**), (b) detached centrosomes (arrow), (c) abnormal PCM morphology (box) and extra DNA (arrow), and (d) cytokinesis failure **(Movie S3)**. (D) Embryos expressing mCherry::histone and mCherry::plasma membrane. Boxed regions highlight cytokinetic furrow shown in bottom panels. Time-lapse recordings of cleavage furrow formation and ingression in L4440 and *kin-3(RNAi)* embryos. (E) CK2 depletion leads to a delay in cell cycle progression: (Top) Measurement of cell cycle lengths (average ± s.d.) from first metaphase to second metaphase in AB or P1 cell. Each dot represents an embryo. (Bottom) Wild-type embryo representing cell cycle stages used for quantification. Note that second metaphase of the anterior blastomere (AB: arrow) initiates before second metaphase of the posterior blastomere (P1: arrow). (A,B,E) Box ranges from the first through third quartile of the data, and thick bar represents the median. Dashed line extends 1.5 times the inter-quartile range or to the minimum and maximum data point. \*\*\**p* <0.001 (two-tailed t-test). Scale bar, 10 μm.

To understand the cause of embryonic lethality by CK2 depletion, we examined early embryos expressing GFP::β-tubulin, mCherry::γ-tubulin and mCherry::histone (Toya et al., 2010) using confocal live imaging (**Fig. 4C, Movie S2, S3**). During mitotic cell division, *kin-3(RNAi)* resulted in various cell cycle defects, including aberrant chromosome segregation (**Fig. 4Ca, Movie S2)**, detached centrosomes where centrosomes lose association with the nucleus (**Fig. 4Cb)**, abnormal PCM morphology (**Fig. 4Cc)**, and cytokinesis failure (**Fig. 4Cd**, **Movie S3**) resulting from incomplete cytokinetic furrow formation (**Fig. 4D, Movie S3)**. We also observed abnormal meiotic divisions, likely leading to extra DNA in embryos **(Fig. S4).** Further, 4D time-lapse imaging revealed a significant cell cycle delay in *kin-3(RNAi)* embryos (**Figs 4E, S5, Movies S2,3**). Together, our results indicate that CK2 is required for proper cell divisions and thus essential for *C. elegans* embryogenesis.

### KIN-3, the catalytic subunit of CK2 exhibits dynamic changes in subcellular localization during the cell cycle

To determine how CK2 might function in the early cell cycle, we examined the subcellular localization of KIN-3 in early embryos. Live imaging of embryos expressing KIN-3::GFP revealed dynamic localization patterns during cell cycle progression (**Fig. 5**). During interphase and early mitosis, KIN-3 is enriched at nuclei. Upon nuclear envelope breakdown (NEBD), KIN-3 appears to localize to the metaphase spindle and remain associated with spindle microtubules and centrosomes throughout mitosis (**Fig. S6)**. During later stages of cell division, KIN-3 is highly enriched in the cytokinetic midbody (**Fig. 5A,B**), which is evidenced by co-localization of KIN-3::GFP and mCherry tagged plasma membrane (Green et al., 2013, Kachur et al., 2008) (**Fig. 5B**). KIN-3::GFP localizes within a spherical area associated with the cytokinetic furrow that is surrounded by the plasma membrane supporting that KIN-3 localizes to the midbody. *kin-3(RNAi)* nearly abolished KIN-3::GFP expression (**Fig. 5A)**, suggesting that KIN-3::GFP expression represent localization patterns specific to KIN-3.

**Figure 5.**
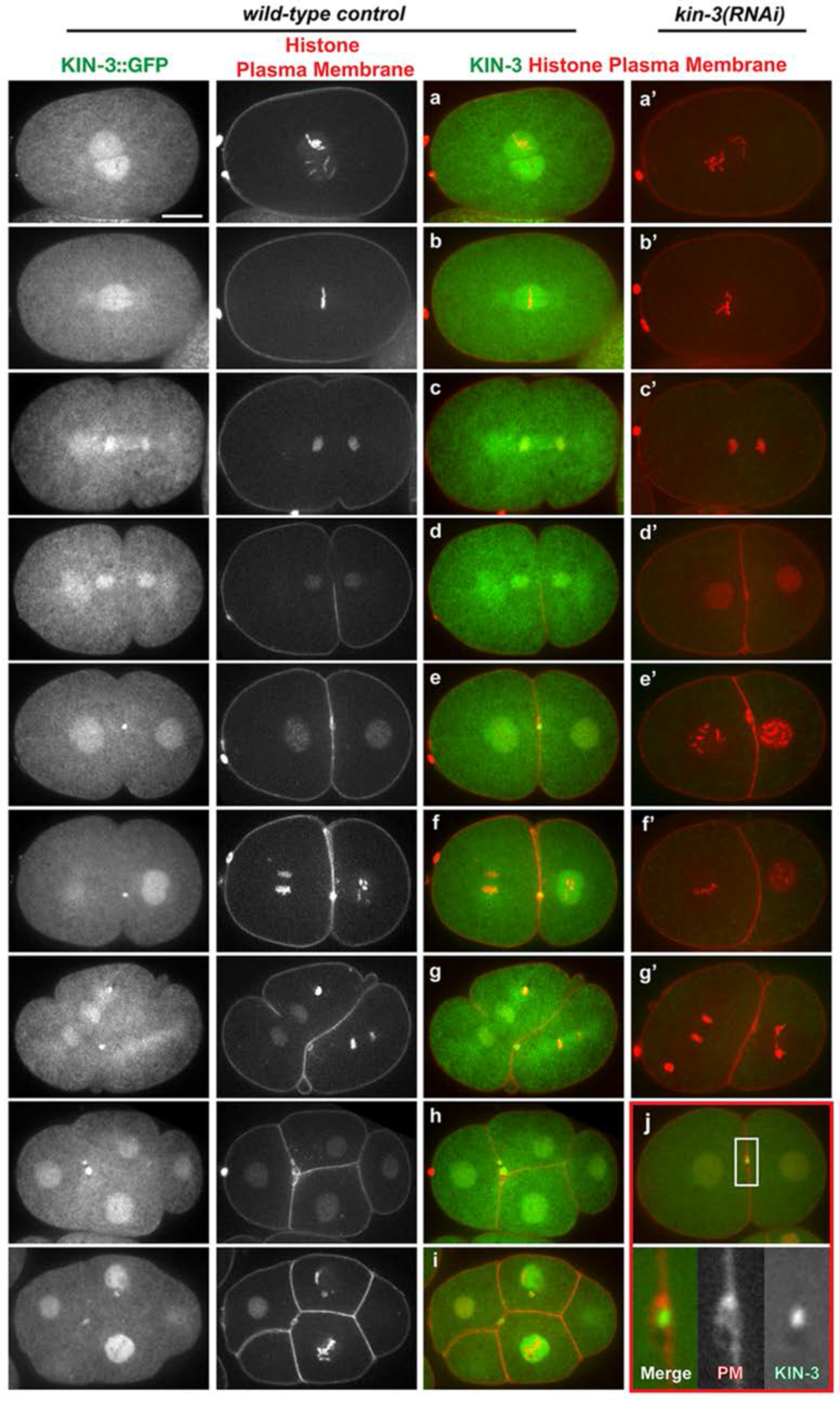
Subcellular localization of KIN-3::GFP. Still images of embryos expressing KIN-3::GFP, mCherry::histone and mCherry::plasma membrane, illustrating that KIN-3 is enriched in nuclei at prophase (a), localizes to the mitotic spindle and centrosomes at mitosis including metaphase (b), anaphase (c) and telophase (d) (see **Fig. S6**). At completion of the first cytokinesis, KIN-3 becomes highly enriched at the midbody-associated structure (e-i, arrows). KIN-3 localization at the nuclei, mitotic spindles and midbody can be observed during second and later cell cycle stages (f-i). All subcellular localizations of KIN-3::GFP are abolished by depletion of KIN-3 by RNAi (a'-g'). The midbody localization of KIN-3::GFP is highlighted (j) by co-localization of plasma membrane (PM). Insets are magnified 4 folds. Scale bar, 10 μm.

Given our finding on KIN-3 localization at the midbody and its related role in cytokinesis, we asked if KIN-3 functions at the midbody to regulate cell division. The Aurora/Ipl1p-related kinase AIR-2 regulates multiple steps of cell division including chromosome segregation and cytokinesis (Schumacher et al., 1998). In particular AIR-2 plays a key role in cytokinesis through ZEN-4, both functioning at the midbody (Kaitna et al., 2000, Severson et al., 2000). To address the possible role of CK2 at the midbody, we tested a genetic interaction between *kin-3* and *air-2* or *zen-4* (**Fig. S7, Table S2**). *kin-3(RNAi)* led to an increase in embryonic lethality of *air-2(or207)* or *zen-4(or153) (* Severson et al., 2000), suggesting that *kin-3* has a positive genetic interaction with *air-2* and *zen-4*, two genes associated with the midbody. Thus, it seems likely that KIN-3 function in early cell cycle as a component of the midbody structure.

### CK2 dependent phosphorylation likely functions in centrosome duplication and the cell cycle

To further support the CK2 holoenzyme-dependent regulation of *zyg-1*, we tested whether chemical inhibition of CK2 could restore embryonic viability to *zyg-1(it25).* We used the highly selective chemical inhibitor of CK2, 4,5,6,7-tetrabromobenzotriazole (TBB) that competes for binding at the ATP-binding site of CK2 (Sarno et al., 2001, Szyszka et al., 1995). It has been reported that TBB specifically abolishes CK2-dependent phosphorylation through *in vitro* kinase assay without affecting the expression levels of CK2 subunits (Alessi et al., 2015, Pagano et al., 2008, Sarno et al., 2001, Szyszka et al., 1995, Wang et al., 2014, Yde et al., 2008). Compared to DMSO controls, TBB treatment partially restores embryonic viability and bipolar spindle formation to *zyg-1(it25)* animals (**Fig. 6A,C,D**). Consistent with CK2 depletion by RNAi, we also observed that TBB-treated embryos possess increased levels of ZYG-1 at centrosomes (1.48 ± 0.57 fold; *p* <0.001) compared to the DMSO control (**Fig. 6E,F**). While TBB produced no significant effect in embryonic viability, TBB treated animals produced a significantly reduced number of progeny (**Fig 6B**) that was observed by RNAi knockdown. Cytological analysis further confirmed that embryos treated with TBB exhibit cell cycle defects including detached centrosomes, DNA missegregation and abnormal PCM morphology (**Fig. 6G**). Taken together, our data suggest that the protein kinase CK2 holoenzyme functions in centrosome duplication and cell division, likely through CK2 holoenzyme-dependent phosphorylation.

**Figure 6.**
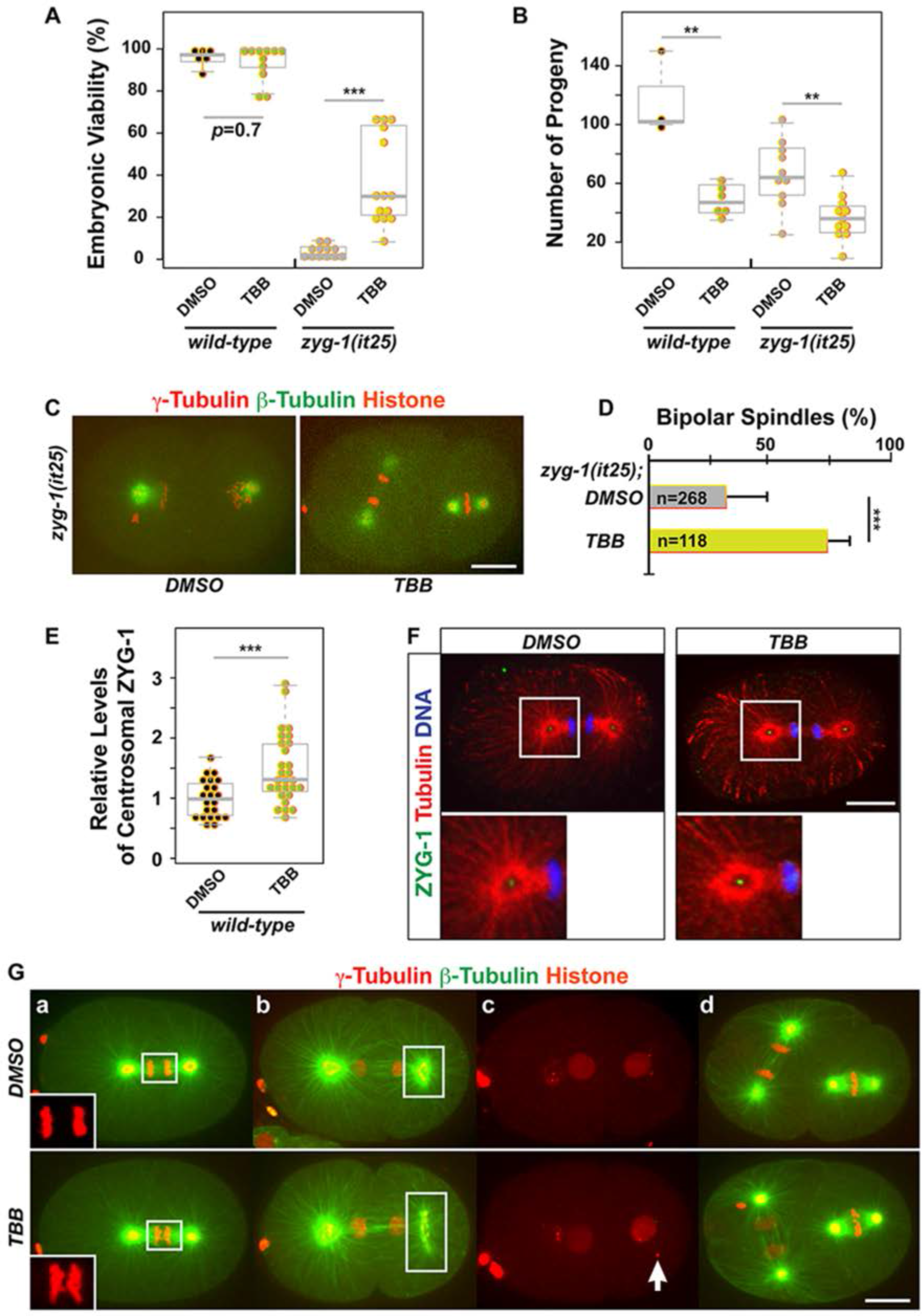
TBB, the chemical inhibitor of CK2, phenocopies CK2 depletion. (A) TBB treatment restores embryonic viability to *zyg-1(it25)* animals at 22.5°C. (DMSO control: 3 ± 3%; TBB: 37 ± 22%), but TBB had a mild effect on wild-type worms (DMSO: 95 ± 6%; TBB: 92 ± 8%). (B) TBB treatment leads to a significant reduction in brood size [wild-type (DMSO: 117 ± 28; TBB: 49 ± 11), *zyg-1(it25)* (DMSO: 67 ± 23; TBB: 39 ± 13)] (A, B) Each dot represents an animal. (C) TBB-treated *zyg-1(it25)* embryo exhibits bipolar spindles, but DMSO control embryo with monopolar spindles. (D) TBB treatment enhances bipolar spindle formation in *zyg-1(it25)* embryos (DMSO: 32 ± 16%; TBB: 73 ± 9.5%, n=blastomeres). Error bars are s.d. (E) Quantification of centrosomal ZYG-1 levels in TBB-treated wild-type embryos. Each dot represents a centrosome. (A,B,E) Box ranges from the first through third quartile of the data. Thick bar represents the median. Dashed line extends 1.5 times the inter-quartile range or to the minimum and maximum data point. (F) TBB-treated wild-type embryo shows more intense ZYG-1 focus at centrosomes. Insets illustrate centrosomal regions magnified 4 folds. (G) Wild-type embryos treated with TBB exhibit defective cell divisions that are similar to RNAi-mediated depletion of CK2: (a) lagging DNA (boxes: Insets are 2 folds), (b) expanded PCM (boxes), (c) detached centrosomes (arrow), (d) cell cycle delay: note cell cycle stages in AB (left) relative to P1 (right) cell. Shown are still images from live confocal imaging. \*\*\**p* <0.001, \*\**p* <0.01 (two-tailed t-test). Scale bar, 10 μm.

## Discussion

### *C. elegans* protein kinase CK2 functions in cell division during embryogenesis

In this study, we have investigated the function of the holoenzyme CK2 in early *C. elegans* embryos. Our data suggest that both the catalytic (KIN-3/CK2α) and regulatory (KIN-10/CK2β) subunits of CK2 are required for the survival of *C. elegans* embryos, which is consistent with prior genome-wide studies (Fraser et al., 2000, Sonnichsen et al., 2005). A fraction of embryos exposed to *kin-3(RNAi)* or *kin-10(RNAi)* were able to successfully complete embryogenesis, although this is likely due to incomplete knockdown of CK2 activity by RNAi feeding. Cytological analyses reveal that RNAi-based depletion of CK2 results in abnormal cell divisions, including chromosome missegregation, a delay in cell cycle progression, cytokinesis failure, as well as aberrant centrosome behaviors. Furthermore, treating embryos with TBB, a chemical inhibitor of CK2, produces similar effects on early cell divisions to those by RNAi knockdown. TBB inhibits CK2 kinase activity via competitive binding to the ATP-binding site of CK2 (Sarno et al., 2001) and it has been reliably utilized for the specific inhibition of CK2 function in the *C. elegans* system (Alessi et al., 2015, Wang et al., 2014). Thus, it seems likely that protein kinase activity of the CK2 holoenzyme is responsible for proper cell division.

The pleiotropic phenotypes caused by CK2 depletion are likely a combined outcome of multiple substrates targeted by CK2 (Meggio and Pinna, 2003). The most prevalent phenotype we observed is abnormal chromosome segregation (**Fig. 4C**, 68%, n=145), including chromosome misalignment at metaphase, lagging DNA at anaphase, and extra DNA during early cell divisions in *C. elegans* embryos. Such roles appear to be evolutionarily conserved in yeast and human cells (Peng et al., 2011, St-Denis et al., 2009). For example, CK2 is known to phosphorylate kinetochore factors (Ndc10, Mif2/CENP-C) in budding yeast (Peng et al., 2011), and the microtubule plus-end tracking protein CLIP-170 that ensures kinetochores attachment to mitotic spindles in human cells (Li et al., 2010). In addition, CK2 targets the chromosome passenger complex (survivin/BIR-1) that functions in chromosome segregation and cytokinesis in mammalian cells (Barrett et al., 2011, Kitagawa and Lee, 2015) and phosphorylates Mad2 to regulate the spindle assembly checkpoint in yeast (Shimada et al., 2009). While we do not know specific cell cycle regulators phosphorylated by *C. elegans* CK2 in early cell division, it is plausible that *C. elegans* CK2 functions in chromosome segregation and cell cycle progression, likely through multiple factors targeted by protein kinase CK2.

The subcellular localization of KIN-3, the catalytic subunit of *C. elegans* CK2, appears to correlate well with the function of CK2 in cell division. Our confocal live imaging reveal that KIN-3::GFP exhibits highly dynamic patterns of subcellular localization during the cell cycle. KIN-3 localizes to nuclei during interphase and early mitosis, and associates with mitotic spindles and centrosomes at mitosis. At later steps of cell division, KIN-3 becomes highly enriched as a distinct focus at the midbody. Our observations are consistent with subcellular localization of the catalytic subunit CK2α described in human cells, where CK2α localizes to nuclei (Penner et al., 1997), midbodies (Salvi et al., 2014), microtubules (Lim et al., 2004) and centrosomes (Faust et al., 2002). In the *C. elegans* germ line, the regulatory subunit KIN-10 is also shown to localize at centrosomes in mitotic cells (Wang et al., 2014). Many known midbody components are known to be required for proper cytokinesis (Green et al., 2012). A role for CK2 at the midbody is further supported by a proteomic survey of the mammalian midbody, where the regulatory subunit CK2β was identified (Skop et al., 2004). Our mass spectrometry analysis also suggests that KIN-3 physically associates with several known midbody proteins including ZEN-4 and CYK-4 (**Table S1**) (Jantsch-Plunger et al., 2000, Raich et al., 1998). In addition, we show that *kin-3* exhibits a positive genetic interaction with *air-2* and *zen-4* that encode midbody proteins regulating cytokinesis (Kaitna et al., 2000, Severson et al., 2000), suggesting that CK2 functions as a part of the midbody structure. Therefore, the holoenzyme CK2 is required for early cell division in *C. elegans* embryos, which appears to be conserved between nematodes and mammals.

### CK2 negatively regulates centrosome duplication in the *C. elegans* embryo

Protein kinase CK2 was identified as part of the SZY-20 immunocomplex. Given that *szy-20* is a genetic suppressor of *zyg-1 (* Kemp et al., 2007, Song et al., 2008), we speculated that CK2 might have a role in centrosome assembly. Our results show that inhibiting either subunit of CK2 restores centrosome duplication and embryonic viability to *zyg-1(it25)* mutants at the semi-restrictive temperature, suggesting the *C. elegans* CK2 holoenzyme functions in centrosome assembly as a negative regulator. Depletion of CK2 does not significantly restore embryonic viability to *zyg-1(it25)* mutants at the restrictive temperature, 24°C, suggesting that loss of CK2 activity does not bypass the requirement for ZYG-1 in early embryos. Furthermore, *kin-3* appears to exhibit a positive genetic interaction with *szy-20* in regulating centrosome duplication, consistent with their physical association. Inhibiting CK2 kinase activity with TBB leads to restoration of centrosome duplication and embryonic viability to *zyg-1(it25)* mutants, indicating that CK2-dependent phosphorylation plays a critical role in centrosome duplication. While it has been shown that aberrant CK2α activity leads to centrosome amplification in mammalian cells (St-Denis et al., 2009), it remains unclear how CK2 function is linked to centrosome assembly. Given that KIN-3 localizes at centrosomes, CK2 might influence centrosome assembly via phosphorylation of centrosome-associated factors, although it is also possible that CK2 targets centrosome regulators in the cytoplasm before they are recruited to centrosomes.

Our work suggests that *C. elegans* CK2 might function in centrosome duplication by targeting ZYG-1. Both RNAi and TBB mediated inhibition of CK2 function led to elevated levels of ZYG-1 at centrosomes, suggesting that CK2-dependent phosphorylation regulates ZYG-1 by controlling either localization or stability. ZYG-1 phosphorylation by CK2 might interfere with ZYG-1 recruitment to centrosomes. Alternatively, CK2-dependent phosphorylation of ZYG-1 may be a targeting signal for proteasomal degradation. Thus, inhibiting CK2 activity prevents ZYG-1 from degradation, increasing overall ZYG-1 abundance and thereby centrosomal levels. The latter hypothesis is intriguing given that phosphorylation is shown to be required for proteosomal degradation of the ZYG-1 homolog Plk4 (Cunha-Ferreira et al., 2013, Guderian et al., 2010, Holland et al., 2010, Klebba et al., 2013). In either way, increased ZYG-1 levels at centrosomes by inhibiting CK2 activity, at least partially, explains how reducing CK2 activity restores centrosome duplication to *zyg-1(it25)* embryos. It has been known that CK2 is a constitutively active Ser/Thr kinase that favors a conserved target motif including acidic amino acid residues near the phosphorylated residue (Salvi et al., 2009). Although it is beyond the scope of our current study, identifying substrates and specific amino acid residues targeted by CK2 will help in understanding how CK2 regulates centrosome duplication, in particular, how protein kinase CK2 influences ZYG-1 levels at centrosomes in *C. elegans* embryos. In any event, our data suggest that the holoenzyme CK2 functions to influence ZYG-1 levels at centrosomes through its kinase activity and thus, we report the protein kinase CK2 as a negative regulator of centrosome duplication.

In this study, we investigated the role of the conserved protein kinase CK2 in early *C. elegans* embryos, and show that CK2 acts as a negative regulator of centriole duplication and is required for proper cell cycle progression and cytokinesis.

## Materials and Methods

### *C. elegans* strains and genetic analysis

The *C. elegans* strains used in this study were maintained on MYOB plates seeded with *E. coli* OP50. All strains were derived from the wild-type Bristol N2 strain using standard genetics (Brenner, 1974, Church et al., 1995). Strains were maintained at 16 or 19°C unless otherwise indicated. A full list of strains used in this study is listed in **Table S3**. The KIN-3::GFP::3xFLAG strain (MTU5) was generated by standard particle bombardment (Praitis et al., 2001). The KIN-3::GFP::3XFLAG construct was acquired from TransgenOme, which covers 22kbp of the *kin-3* 5'UTR and 9kbp of the *kin-3* 3'UTR (Sarov et al., 2012). RNAi feeding was performed as previously described, and the L4440 empty feeding vector was used as a negative control (Kamath et al., 2003).

For embryonic viability and brood size assays, individual L4 animals were transferred to new plates and allowed to self-fertilize for 24 hours at the temperatures indicated. For extended RNAi treatments (36-48hours), animals were transferred to a new plate in 24 hours, and allowed to self-fertilize for an additional 24 hours before removal. Progeny were allowed at least 24 hours to complete embryogenesis before counting the number of hatched larvae and unhatched (dead) eggs.

### Cytological Analysis

For immunostaining, the following primary antibodies were used at 1:2,000-3,000 dilutions: α-Tubulin: DM1a (Sigma, St-Louis, MO, USA), α-GFP: IgG1κ (Roche, Indianapolis, IN, USA), α-ZYG-1 (Stubenvoll et al., 2016), α-TBG-1 (Stubenvoll et al., 2016), α-SAS-4 (Song et al., 2008), α-SAS-5 (this study; **Fig. S8B**) and α-SZY-20 (Song et al., 2008). Alexa Fluor 488 and 561 (Invitrogen, Carlsbad, CA, USA) were used as secondary antibodies. Affinity-purified rabbit polyclonal antibody for KIN-3 was generated (YenZym, South San Franscisco, CA, USA) against the following peptides (aa361-373): Ac-VEDSSDHEEDVVV-amide. Immunostaining was performed as described previously (Song et al., 2008).

Confocal microscopy was performed as described in (Stubenvoll et al., 2016) using a Nikon Eclipse Ti-U microscope equipped with a Plan Apo 60 x 1.4 NA lens, a Spinning Disk Confocal (CSU X1) and a Photometrics Evolve 512 camera. Images were acquired using MetaMorph software (Molecular Devices, Sunnyvale, CA, USA). MetaMorph was used to draw and quantify regions of fluorescence intensity and Adobe Photoshop CS6 was used for image processing. To quantify centrosomal signals (SPD-2::GFP, TBG-1), the average intensity within a 25-pixel (1 pixel = 0.151 μm) diameter region was measured within an area centered on the centrosome and the focal plane with the highest average intensity (corresponding to the centrosome) was recorded. The average fluorescence intensity within a 25-pixel diameter region drawn outside of the embryo was used for background subtraction. Centriolar signals (ZYG-1, SAS-5, SAS-4) were quantified in the same manner, except that 8-pixel diameter regions were used.

### Immunoprecipitation

Embryos were extracted by bleaching gravid worms in a hypochlorite solution [1:2:1 ratio of M9 buffer, bleach (5.25% sodium hypochlorite) and 5M NaCl], washed with M9 buffer, flash-frozen in liquid nitrogen and stored at -80°C until use. For α-GFP immunoprecipitation experiments, 20 μL of Mouse-α-GFP magnetic beads (MBL, Naka-ku, Nagoya, Japan) were used per reaction. Beads were prepared by washing twice for 15 minutes in PBST (PBS; 0.1% Triton-X), followed by a third wash in 1x lysis buffer (50 mM HEPES [pH 7.4], 1mM EDTA, 1mM MgCl2, 200 mM KCl, and 10% glycerol (v/v)) (Cheeseman et al., 2004). Embryos were ground in microcentrifuge tubes containing an equal amount of 1x lysis buffer supplemented with complete protease inhibitor cocktail (Roche, Indianapolis, IN, USA) and MG132 (Tocris, Avonmouth, Bristol, UK) and briefly sonicated prior to centrifugation. Samples were spun in a desktop centrifuge twice for 20 minutes, collecting the supernatant after each spin. Protein quantification was then determined using a NanoDrop spectrophotometer (Thermo-Fisher, Hanover Park, IL, USA) and adjusted such that the same amount of total protein was used for each reaction. Beads were then added to the microcentrifuge tubes containing embryonic lysates. Samples were incubated and rotated for one hour at 4°C and subsequently washed three times with PBST. Samples were then resuspended in 20 μL of a solution containing 2X Laemmli Sample Buffer (Sigma, St-Louis, MO, USA) and 10% β-mercaptoethanol (v/v) and boiled for five minutes. Mass spectrometry analysis was performed as described previously (Song et al., 2011).

### Western Blotting

For western blotting experiments, samples were sonicated for 5 minutes and boiled in a solution of 2X Laemmli Sample Buffer and 10% β-mercaptoethanol before being fractionated on a 4-12% NuPAGE Bis-Tris gel (Invitrogen, Carlsbad, CA, USA). The iBlot Gel Transfer system (Invitrogen, Carlsbad, CA, USA) was then used to transfer samples to a nitrocellulose membrane. The following antibodies were used at 1:3,000-10,000 dilutions: α-Tubulin: DM1a (Sigma, St-Louis, MO, USA), α-GFP: IgG1κ (Roche, Indianapolis, IN, USA), α-SPD-2 (Song et al., 2008), α-SZY-20 (Song et al., 2008), α-TBG-1 (Stubenvoll et al., 2016), α-SAS-5 (Song et al., 2011) and α-KIN-3 (this study; **Fig. S8A**). IRDye secondary antibodies (LI-COR Biosciences, Lincoln, NE, USA) were used at a 1:10,000 dilution. Blots were imaged using the Odyssey infrared scanner (LI-COR Biosciences, Lincoln, NE, USA), and analyzed using Image Studio software (LI-COR Biosciences, Lincoln, NE, USA). Affinity-purified rabbit polyclonal antibody for KIN-3 was generated ((YenZym, South San Franscisco, CA, USA) against the following peptides (aa338-356): Ac-CQADGQGASNSASSQSSDAK-amide.

### TBB treatment

MYOB plates were first seeded with OP50 bacteria and allowed to dry overnight. The media was then supplemented with 0.5 mM TBB (Tocris, Avonmouth, Bristol, UK) dissolved in a solution of 50% DMSO. TBB was added to the surface of plates, such that the final concentration of TBB was 15 μM based on the volume of media, and allowed to soak and diffuse through media overnight. Final TBB concentrations were derived from (Wang et al., 2014). An equal volume of solution containing 50% DMSO was added to plates and used as a control.

### Statistical Analysis

All *p* -values were calculated using two-tailed t-tests assuming equal variance among sample groups. Statistics are presented as average ± s.d. unless otherwise specified. Data were independently replicated at least three times for all experiments and subsequently analyzed for statistical significance.

## Acknowledgements

We are grateful for the assistance of Mass Spectrometry Facility at Harvard Medical School and National Institutes of Health. We also thank Song lab members for experimental assistance and Kevin O'Connell for RNAi and worm stains. Some strains were provided by the CGC, which is funded by NIH Office of Research Infrastructure Programs (P40 OD010440).

## Competing Interests

No competing interests declared.

## Author Contributions

J.C.M. and M.H.S. designed the experiments and wrote the manuscript. J.C.M. and M.H.S. performed quantifications of confocal imaging and protein levels from western blots. J.C.M., M.M.K., M.D.S., L.E.D. and M.H.S. performed experiments and provided data.

## Funding

This work was supported by a grant [7R15GM11016-02 to M.H.S.] from the National Institute of General Medical Sciences, and Research Excellence Fund (to M.H.S) from the Center for Biomedical Research at Oakland University. The funders had no role in study design, data collection and analysis, decision to publish, or preparation of the manuscript.

## Supplementary Information

**Figure S1.**
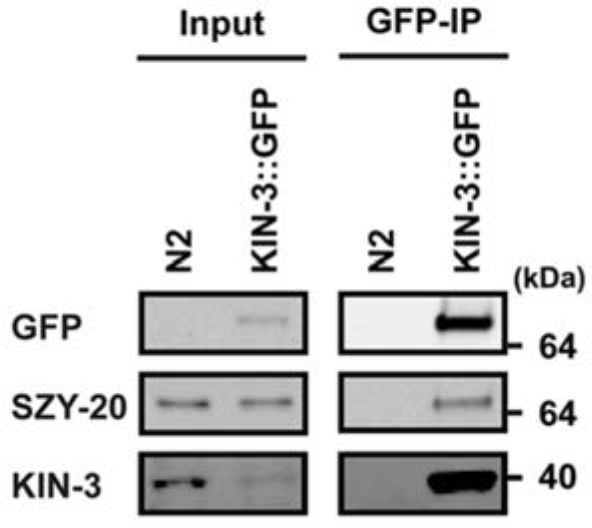
KIN-3/CK2α, the catalytic subunit of CK2, physically interacts with SZY-20 in *C. elegans* embryos. SZY-20 co-precipitates with KIN-3::GFP. Protein lysates from wild-type (*N2*) embryos were used as a negative control. ~5% of total embryonic lysates were loaded in input lanes. While endogenous SZY-20 levels are similar between two samples in input lanes, endogenous KIN-3 levels are much lower in embryos expressing KIN-3::GFP compared to KIN-3 levels in wild-type (*N2*), suggesting that the transgene expression of KIN-3::GFP is likely to down-regulate endogenous KIN-3 expression. However, endogenous KIN-3 co-precipitates with KIN-3::GFP (IP lane), suggesting KIN-3::GFP physically interacts with endogenous KIN-3.

**Figure S2.**
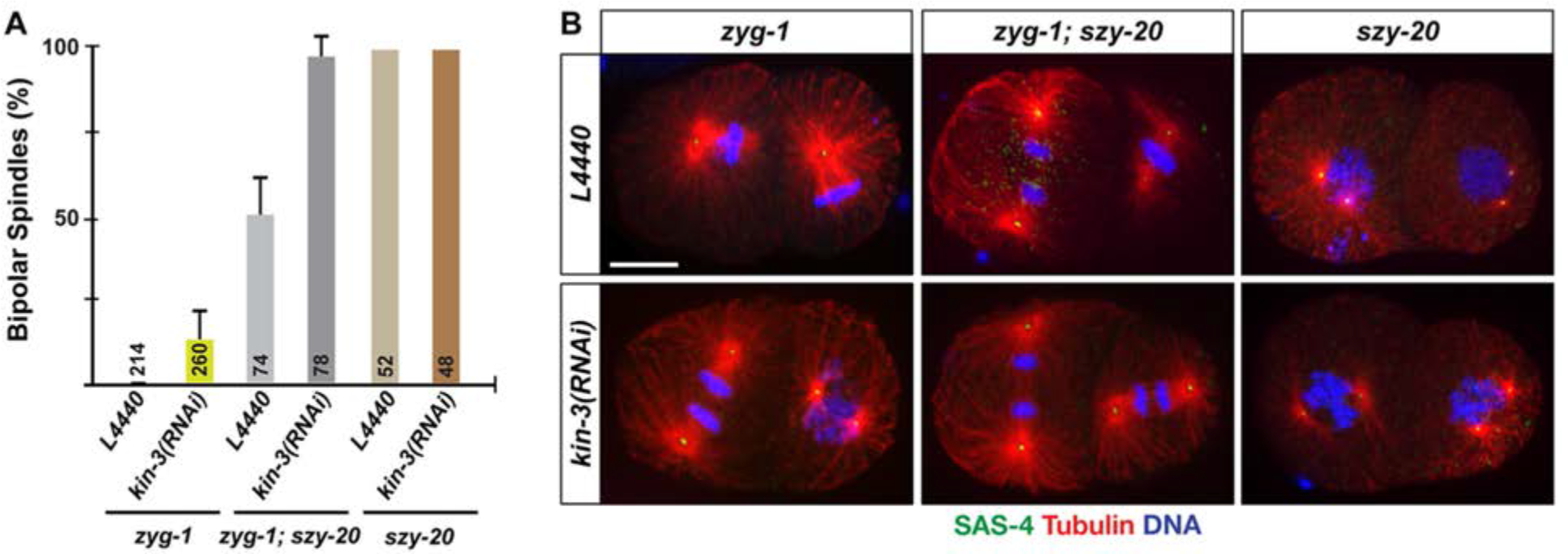
Co-depleting KIN-3 and SZY-20 further enhances bipolar spindle formations to *zyg-1(it25)* embryos. (A) Quantification of bipolar spindle formation in *zyg-1(it25), zyg-1(it25); szy-20(bs52)* and *szy-20(bs52)* embryos treated with *kin-3(RNAi)* and control RNAi (*L4440)* at 24°C, the restrictive temperature for *zyg-1(it25). kin-3(RNAi)* in *zyg-1(it25);szy-20(bs52)* double mutants further increases (95.8 ± 5.9%) bipolar spindle formation, compared to a partial restoration of bipolar spindle formation to *zyg-1(it25)* by *kin-3(RNAi)* (12.9 ± 8.4%) or *szy-20(bs52)* (49.3 ± 11.5%) alone. Average values are shown. Error bars are s.d. n is given as the number of blastomeres scored. (B) Immunostained embryos illustrate monopolar or bipolar spindles at the second mitosis. Scale bar, 10 μm.

**Figure S3.**
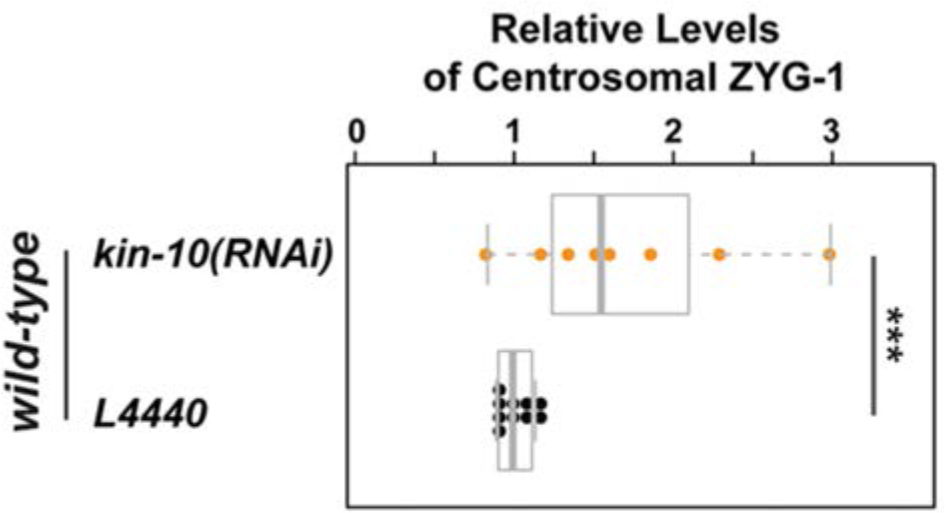
Knocking down KIN-10/CK2β, the regulatory subunit of CK2, leads to increased ZYG-1 levels at centrosomes. Centrosomal ZYG-1 levels at first anaphase were quantified in wild-type embryos treated with *kin-10(RNAi)* or control (*L4440)*. Values are relative to centrosomal ZYG-1 levels in wild-type embryos treated with control. *kin-10(RNAi)* results in an increase in centrosomal ZYG-1 (1.70 ± 0.41 fold). Each dot represents a centrosome. Box ranges from the first through third quartile of the data, and thick bar represents the median. Dashed line extends 1.5 times the inter-quartile range or to the minimum and maximum data point. \*\*\**p* <0.001 (two-tailed t-test).

**Figure S4.**
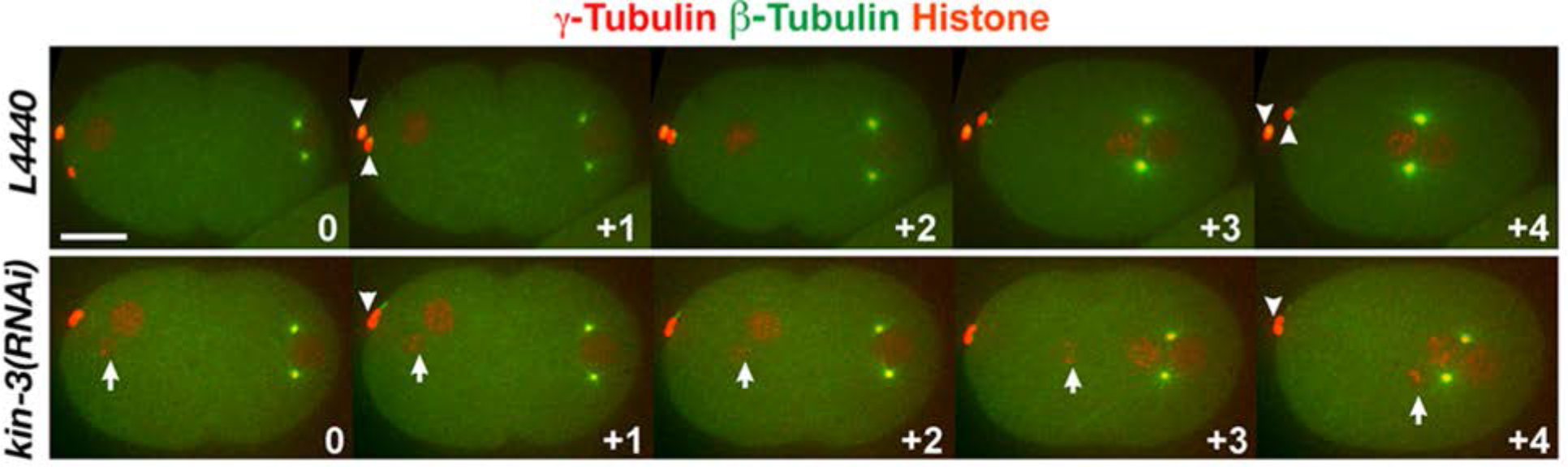
Defective meiotic divisions are observed in *kin-3(RNAi).* Shown are still images selected from 60 sec-interval 4D time-lapse movies. *kin-3(RNAi)* results in an error in meiotic division, likely leading to extra DNA (arrows). While control embryo exhibits two polar bodies (arrowheads), *kin-3(RNAi)* embryo shows only one polar body. Although we cannot rule out the possibility that the second polar body is out of focus in this movie, it is also possible that KIN-3 depletion may lead to polar body extrusion failure and extra DNA. Time is given relative to the start of the movie (end of the second meiotic division). Scale bar, 10 μm.

**Figure S5.**
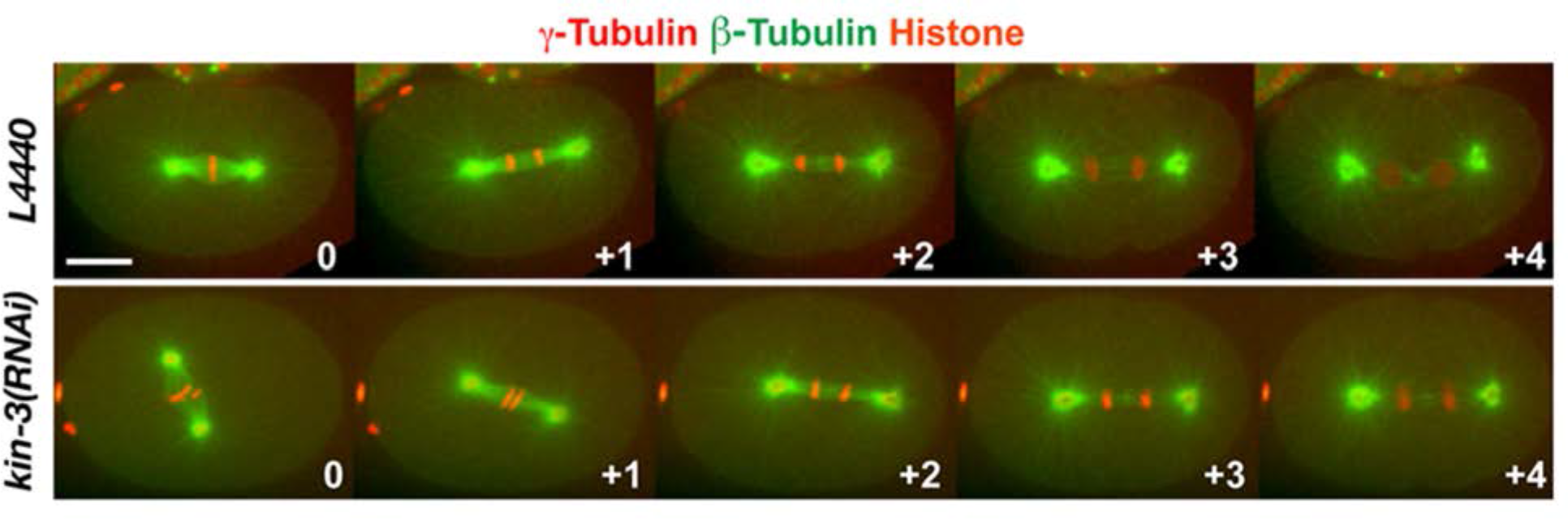
*kin-3(RNAi)* leads to a delay in cell cycle progression. Shown are still images selected from 60 sec-interval 4D time-lapse movies of *kin-3(RNAi)* (n=48 embryos) or control RNAi (n=42 embryos). While the control embryo progresses from metaphase (t=0) to telophase (t=+4) during the shown time interval, the *kin-3(RNAi)* embryo progresses from metaphase (t=0) to anaphase (t=+4) during the same time interval. Time (min) is relative to the first metaphase. Scale bar, 10 μm.

**Figure S6.**
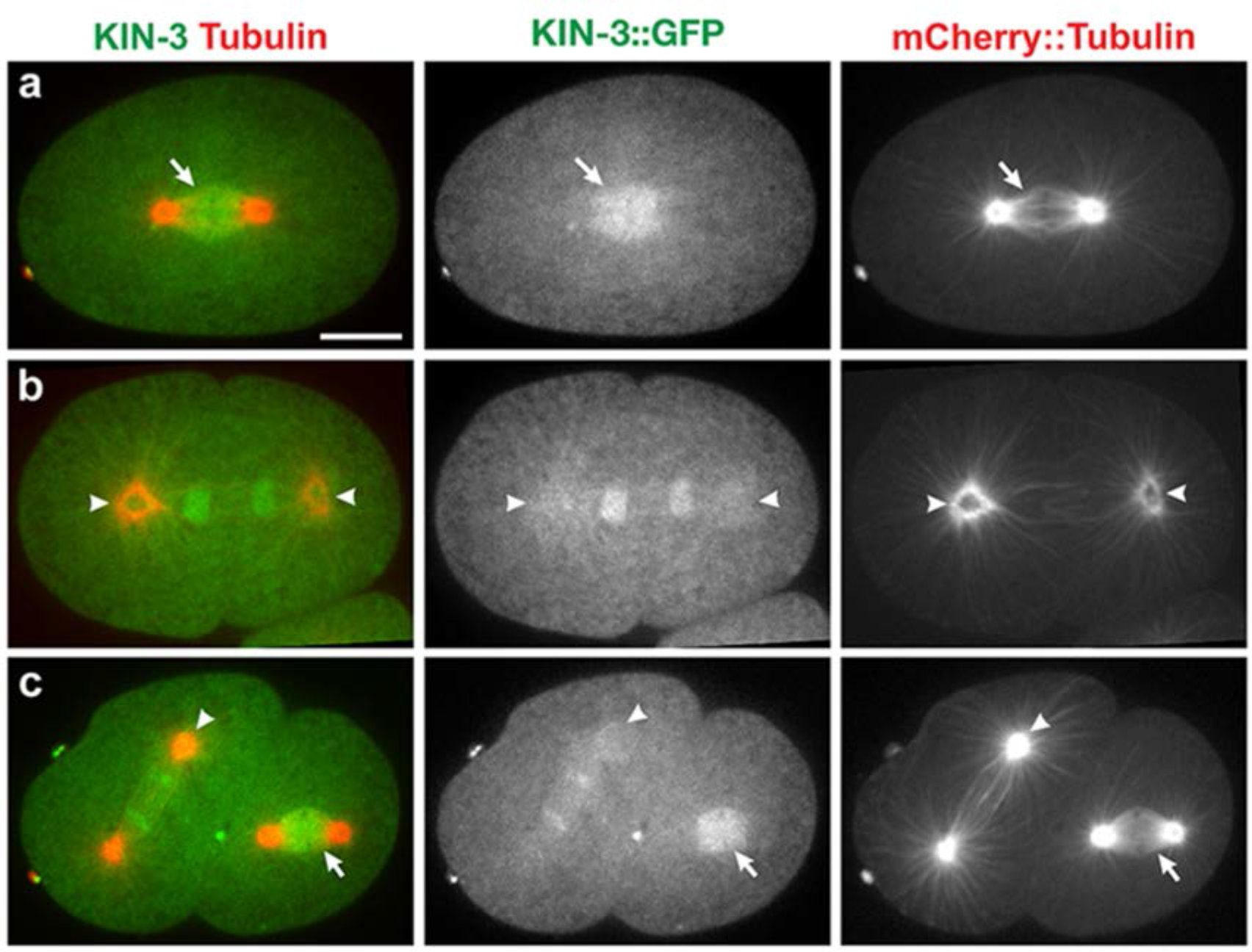
KIN-3 localizes to the mitotic spindle and centrosomes. KIN-3::GFP co-localizes with mCherry::Tubulin at spindle microtubules (arrows) at the first (a) and second (c) metaphase. During late mitosis in the first (b) and second (c) cell division, KIN-3::GFP appears to be associated with centrosomes (arrowheads) that are highlighted with mCherry::Tubulin co-localization. Shown are still images selected from 4D time-lapse movies. Scale bar, 10 μm.

**Figure S7.**
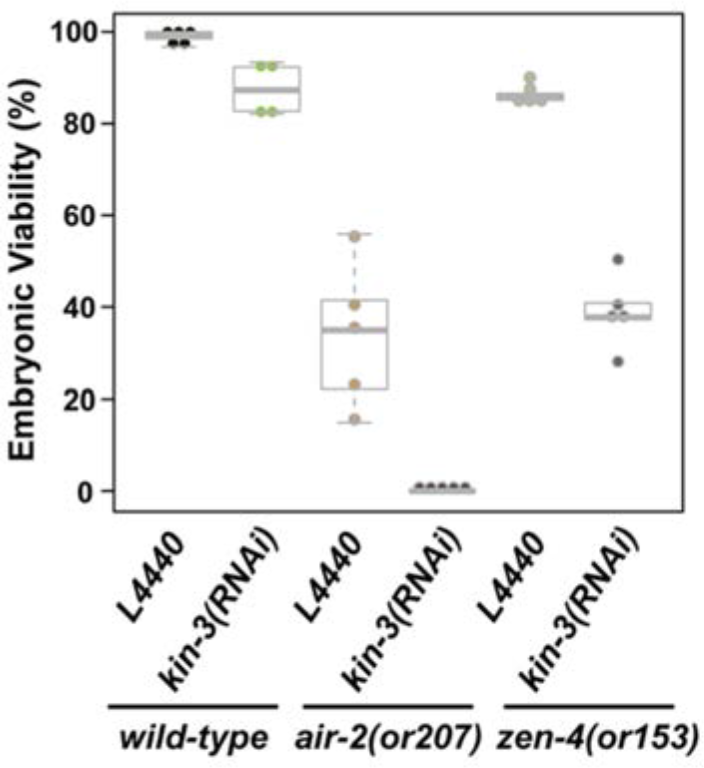
Depleting KIN-3 further decreases embryonic viability in *air-2(or207)* and *zen-4(or153)* mutants. At 17°C, *kin-3(RNAi)* in wild-type worms (n=605) produces 87.5% (± 5.6) embryonic viability and *air-2(or207ts)* mutants (n=672) exhibit 33.0% (± 16.0) embryonic viability. In contrast, *kin-3(RNAi)* in *air-2(or207)* mutants (n=819) leads to 0.3% (± 0.6) embryonic viability, which suggests a positive genetic interaction between *kin-3* and *air-2.* Similarly, *zen-4(or153)* mutants (n=740) exhibit 86.7% (± 2.4) embryonic viability, whereas *kin-3(RNAi)* in *zen-4(or153)* mutants (n=797) leads to 39.0% (± 8.4) embryonic viability at 17°C, showing decreased embryonic viability by co-depleting *kin-3* and *zen-4*. Each dot marks % embryonic viability of the progeny produced by a single worm (n=the total number of progeny scored). Boxes range from the first through third quartile of the data and thick bars represent the median. Dashed lines extend 1.5 times the inter-quartile range, or to the minimum and maximum data point.

**Figure S8.**
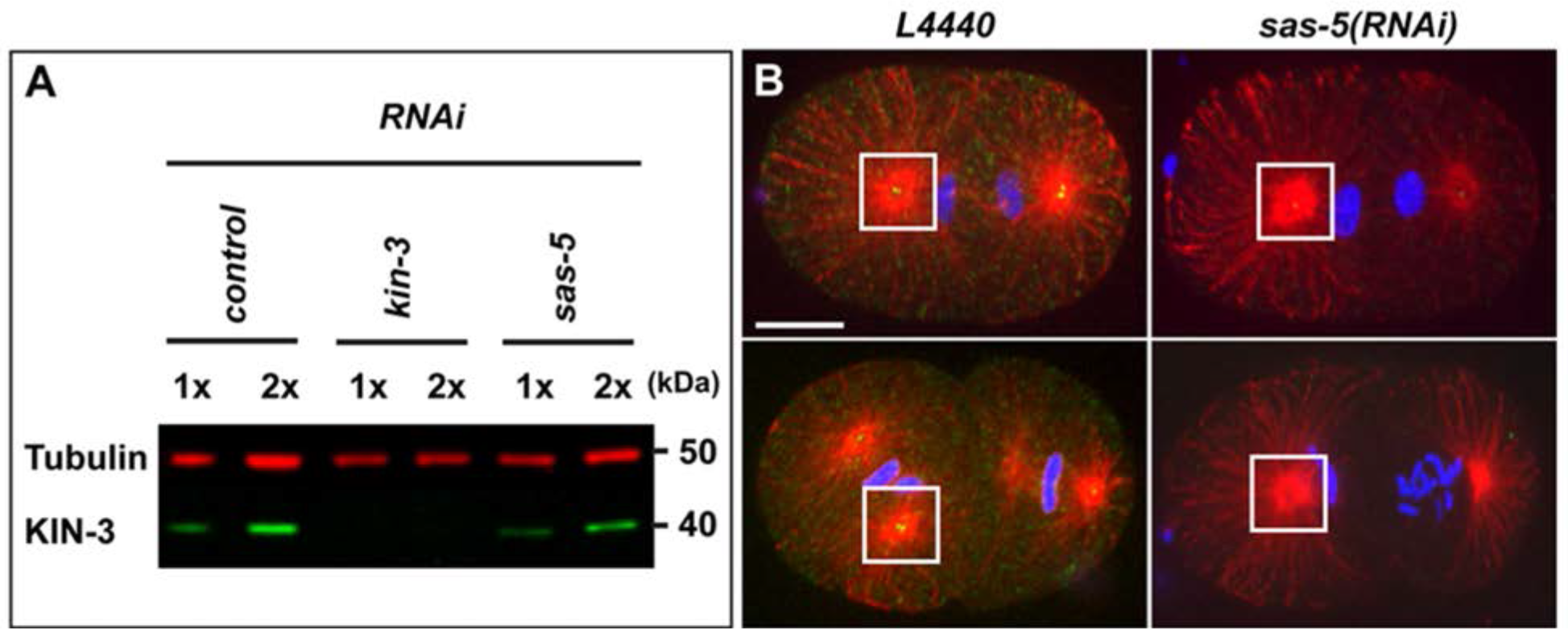
Antibody specificity. (A) Immunoblot illustrates that anti-KIN-3 detects endogenous KIN-3 in control and *sas-5(RNAi)* embryos but detection is abolished by *kin-3(RNAi)*, suggesting the specificity of KIN-3 antibody. Tubulin was used as a loading control. (B) SAS-5 antibodies detect SAS-5 at centrosomes and cytoplasm. However, *sas-5(RNAi)* significantly diminishes SAS-5 detection, indicating the antibody specificity. Boxes highlight centrosomes. Scale bar, 10 μm.

## Supplementary Movies

**Movie S1. Knocking down KIN-3 leads to increased ZYG-1 at centrosomes.** Embryos expressing GFP::ZYG-1-C-term illustrate that *kin-3(RNAi)* leads to elevated levels of centrosome-associated ZYG-1 during the first cell cycle, compared to L4440 controls. Movies are taken 60 sec interval. Time is relative to the first metaphase (t=0) and shown in hh:mm. 5 frames per second (fps). Scale bar, 5 μm.

**Movie S2. KIN-3 is required for proper cell division.** 4D time-lapse movies of embryos expressing mCherry::plasma membrane and mCherry::histone. Compared to control (L4440), *kin-3(RNAi)* results in abnormal chromosome segregation and delay in cell cycle progression. Movies are taken 60 sec interval. Time is relative to the first metaphase (t=0) and shown in hh:mm. 5 fps. Scale bar, 5 μm.

**Movie S3. KIN-3 is required for proper cytokinesis.** 4D time-lapse movies of embryos expressing GFP::tubulin, mCherry::TBG-1 and mCherry::histone. *kin-3(RNAi)* results in a cytokinesis failure following successful centrosome duplication, leading to tetrapolar spindles at the second mitosis. Movies are taken 60 sec interval. Time is relative to the first metaphase (t=0) and shown in hh:mm. 5 fps. Scale bar, 5 μm.

**Movie S4. Subcellular localization of KIN-3::GFP.** Embryo expressing KIN-3::GFP, mCherry::plasma membrane and mCherry::histone illustrates dynamic localization patterns of KIN-3. KIN-3::GFP localizes to nuclei, mitotic spindles and is enriched at the midbody. Movies are taken 60 sec interval. Time is relative to the first metaphase (t=0) and shown in hh:mm. 5 fps. Scale bar, 5 μm.

**Table S1.**
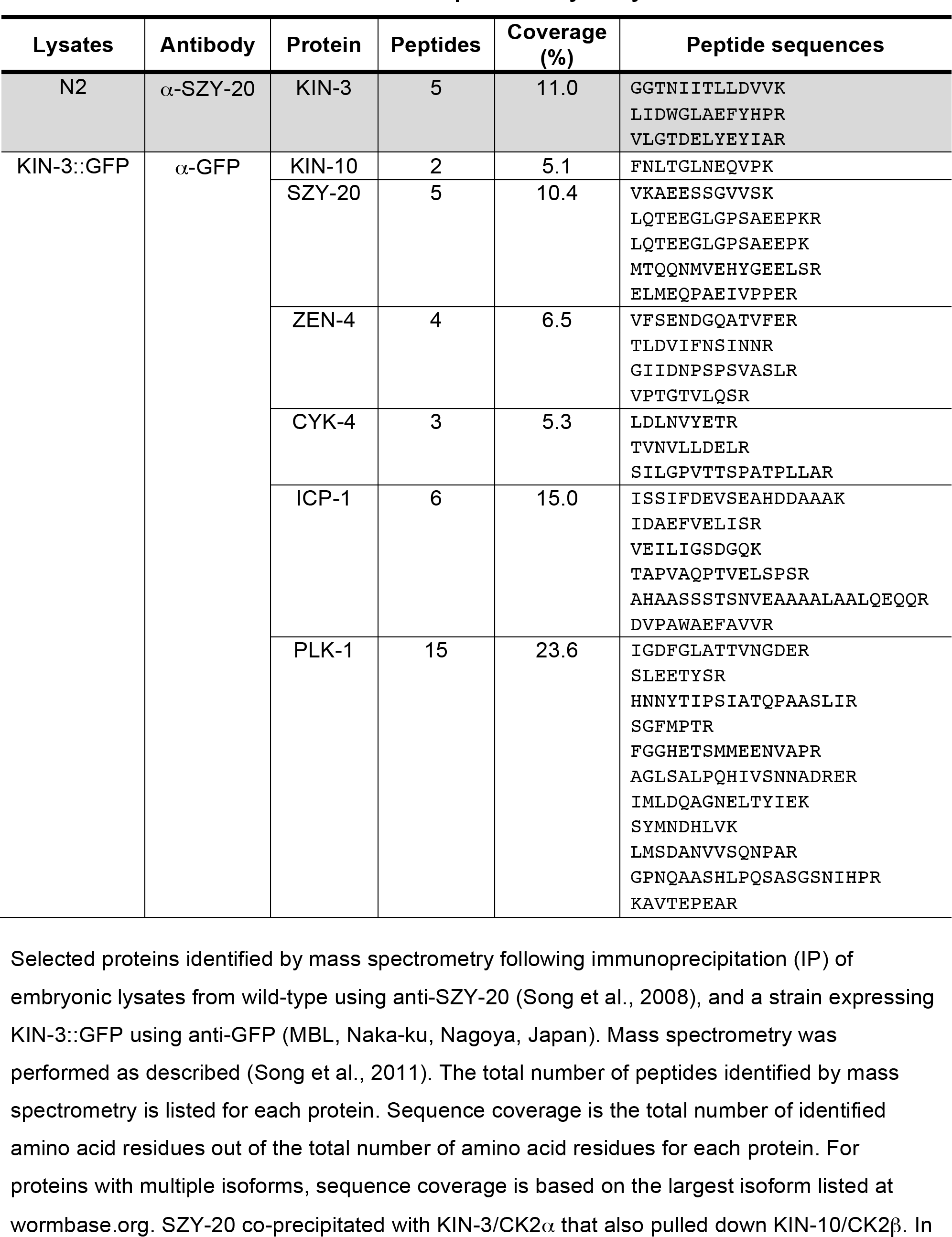
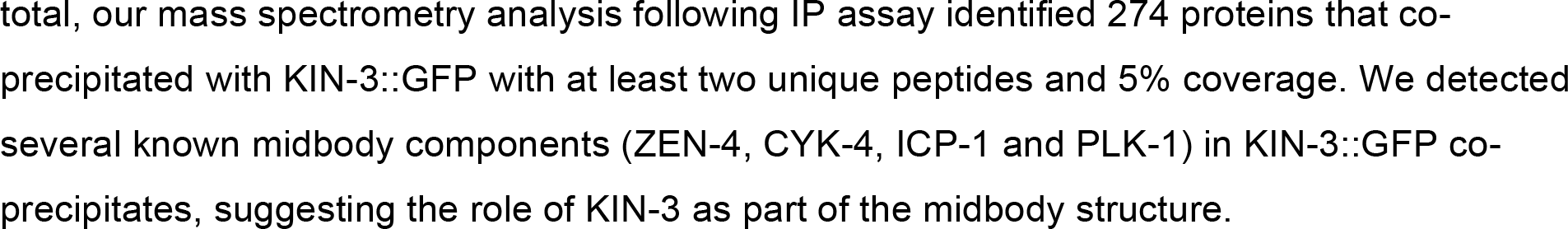
Mass spectrometry analysis.

**Table S2.** Genetic Analysis of CK2.

|  | °C | % Embryonic Viability<br>(ave ± s.d.) | n |
| --- | --- | --- | --- |
| *N2*; L4440 | 24.0<sup>a</sup> | 98.5 ± 1.2 | 3190 |
| *N2*; *kin-3*(RNAi) |  | 55.7 ± 16.1 | 3724 |
| *N2*; *kin-10*(RNAi) |  | 77.4 ± 7.0 | 1147 |
| *zyg-1(it25)*; L4440 |  | 0.2 ± 1.0 | 1090 |
| *zyg-1(it25)*; *kin-3*(RNAi) |  | 0.4 ± 1.2 | 1078 |
| *zyg-1(it25)*; *kin-10*(RNAi) |  | 0.6 ± 1.0 | 576 |
| *szy-20(bs52)*; L4440 |  | 29.1 ± 22.3 | 163 |
| *szy-20(bs52)*; *kin-3*(RNAi) |  | 27.6 ± 15.8 | 238 |
| *zyg-1(it25)*; *szy-20(bs52)*; L4440 |  | 17.8 ± 24.6 | 183 |
| *zyg-1(it25)*; *szy-20(bs52)*; *kin-3*(RNAi) |  | 13.9 ± 20.1 | 382 |
| *N2*; L4440 | 22.5<sup>a</sup> | 98.5 ± 1.4 | 2921 |
| *N2*; *kin-3*(RNAi) |  | 52.8 ± 10.4 | 2433 |
| *N2*; *kin-10*(RNAi) |  | 55.0 ± 20.5 | 1268 |
| *zyg-1(it25)*; L4440 |  | 8.0 ± 12.6 | 7257 |
| *zyg-1(it25)*; *kin-3*(RNAi) |  | 24.8 ± 24.9 | 7389 |
| *zyg-1(it25)*; *kin-10*(RNAi) |  | 40.3 ± 26.1 | 2786 |
| *N2*; DMSO | 22.5<sup>b</sup> | 96.2 ± 2.0 | 276 |
| *N2*; 15μM TBB |  | 96.9 ± 8.1 | 174 |
| *zyg-1(it25)*; DMSO |  | 3.1 ± 3.1 | 637 |
| *zyg-1(it25)*; 15μM TBB |  | 37.0 ± 21.8 | 330 |
| *N2*; L4440 | 17.0<sup>c</sup> | 98.9 ± 1.3 | 702 |
| *N2*; *kin-3*(RNAi) |  | 87.5 ± 5.6 | 605 |
| *air-2(or207)*; L4440 |  | 33.8 ± 16.1 | 672 |
| *air-2(or207)*; *kin-3*(RNAi) |  | 0.3 ± 0.6 | 819 |
| *zen-4(or153)*; L4440 |  | 86.7 ± 2.4 | 740 |
| *zen-4(or153)*; *kin-3*(RNAi) |  | 38.8 ± 8.4 | 797 |

<sup>a</sup>: Presented in Fig. 1
<sup>b</sup>: Presented in Fig. 6
<sup>c</sup>: Presented in Fig. S5

**Table S3.**
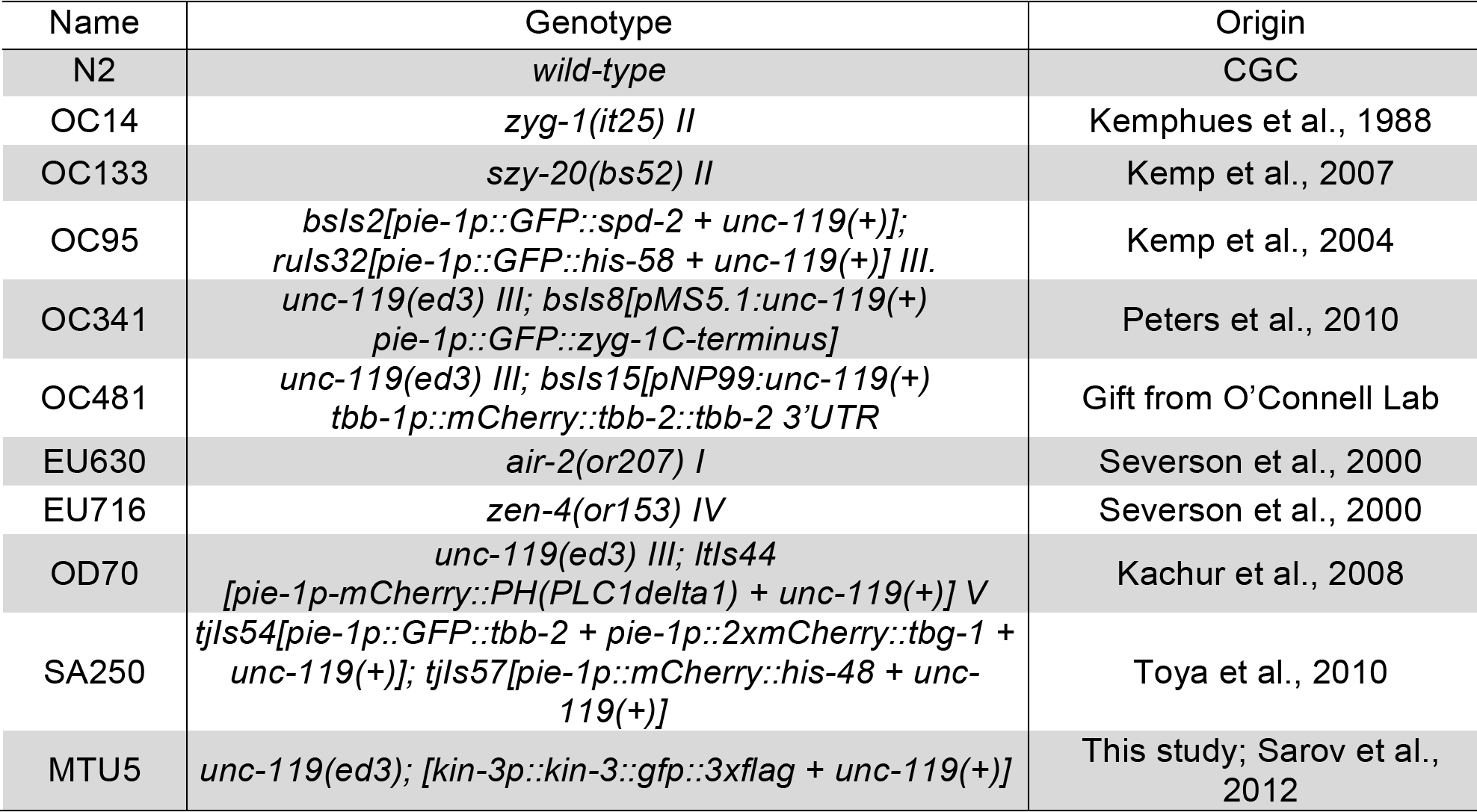
List of *C. elegans s* trains used in this study.

## References

Alessi, A. F., Khivansara, V., Han, T., Freeberg, M. A., Moresco, J. J., Tu, P. G., Montoye, E., Yates, J. R., 3rd, Karp, X. & Kim, J. K. (2015). Casein kinase II promotes target silencing by miRISC through direct phosphorylation of the DEAD-box RNA helicase CGH-1. Proc Natl Acad Sci U S A, 112, E7213–22.

Arquint, C. & Nigg, E. A. (2014). STIL microcephaly mutations interfere with APC/C-mediated degradation and cause centriole amplification. Curr Biol, 24, 351–60.

Barrett, R. M., Colnaghi, R. & Wheatley, S. P. (2011). Threonine 48 in the BIR domain of survivin is critical to its mitotic and anti-apoptotic activities and can be phosphorylated by CK2 in vitro. Cell Cycle, 10, 538–48.

Brenner, S. (1974). The genetics of *Caenorhabditis elegans*. Genetics, 77, 71–94.

Brownlee, C. W., Klebba, J. E., Buster, D. W. & Rogers, G. C. (2011). The Protein Phosphatase 2A regulatory subunit Twins stabilizes Plk4 to induce centriole amplification. J Cell Biol, 195, 231–43.

Cheeseman, I. M., Niessen, S., Anderson, S., Hyndman, F., Yates, J. R., 3rd, Oegema, K. & Desai, A. (2004). A conserved protein network controls assembly of the outer kinetochore and its ability to sustain tension. Genes Dev, 18, 2255–68.

Church, D. L., Guan, K. L. & Lambie, E. J. (1995). Three genes of the MAP kinase cascade, *mek-2, mpk-1/sur-1* and *let-60* ras, are required for meiotic cell cycle progression in *Caenorhabditis elegans*. Development, 121, 2525–35.

Cunha-Ferreira, I., Bento, I., Pimenta-Marques, A., Jana, S. C., Lince-Faria, M., Duarte, P., Borrego-Pinto, J., Gilberto, S., Amado, T., Brito, D., et al. (2013). Regulation of autophosphorylation controls PLK4 self-destruction and centriole number. Curr Biol, 23, 2245–54.

De Groot, R. E., Rappel, S. B., Lorenowicz, M. J. & Korswagen, H. C. (2014). Protein kinase CK2 is required for Wntless internalization and Wnt secretion. Cell Signal, 26, 2601–5.

Decker, M., Jaensch, S., Pozniakovsky, A., Zinke, A., O’connell, K. F., Zachariae, W., Myers, E. & Hyman, A. A. (2011). Limiting amounts of centrosome material set centrosome size in *C. elegans* embryos. Curr Biol, 21, 1259–67.

Delattre, M., Canard, C. & Gonczy, P. (2006). Sequential protein recruitment in *C. elegans* centriole formation. Curr Biol, 16, 1844–9.

Delattre, M., Leidel, S., Wani, K., Baumer, K., Bamat, J., Schnabel, H., Feichtinger, R., Schnabel, R. & Gonczy, P. (2004). Centriolar SAS-5 is required for centrosome duplication in *C. elegans*. Nat Cell Biol, 6, 656–64.

Dzhindzhev, N. S., Tzolovsky, G., Lipinszki, Z., Schneider, S., Lattao, R., Fu, J., Debski, J., Dadlez, M. & Glover, D. M. (2014). Plk4 phosphorylates Ana2 to trigger Sas6 recruitment and procentriole formation. Curr Biol, 24, 2526–32.

Faust, M., Gunther, J., Morgenstern, E., Montenarh, M. & Gotz, C. (2002). Specific localization of the catalytic subunits of protein kinase CK2 at the centrosomes. Cell Mol Life Sci, 59, 2155–64.

Fraser, A. G., Kamath, R. S., Zipperlen, P., Martinez-Campos, M., Sohrmann, M. & Ahringer, J. (2000). Functional genomic analysis of *C. elegans* chromosome I by systematic RNA interference. Nature, 408, 325–30.

Godinho, S. A. & Pellman, D. (2014). Causes and consequences of centrosome abnormalities in cancer. Philos Trans R Soc Lond B Biol Sci, 369.

Gonczy, P. (2015). Centrosomes and cancer: revisiting a long-standing relationship. Nat Rev Cancer, 15, 639–52.

Green, R. A., Mayers, J. R., Wang, S., Lewellyn, L., Desai, A., Audhya, A. & Oegema, K. (2013). The midbody ring scaffolds the abscission machinery in the absence of midbody microtubules. J Cell Biol, 203, 505–20.

Green, R. A., Paluch, E. & Oegema, K. (2012). Cytokinesis in animal cells. Annu Rev Cell Dev Biol, 28, 29–58.

Guderian, G., Westendorf, J., Uldschmid, A. & Nigg, E. A. (2010). Plk4 trans-autophosphorylation regulates centriole number by controlling betaTrCP-mediated degradation. J Cell Sci, 123, 2163–9.

Guerra, B. & Issinger, O. G. (2008). Protein kinase CK2 in human diseases. Curr Med Chem, 15, 1870–86.

Hannak, E., Kirkham, M., Hyman, A. A. & Oegema, K. (2001). Aurora-A kinase is required for centrosome maturation in *Caenorhabditis elegans*. J Cell Biol, 155, 1109–16.

Hinchcliffe, E. H., Li, C., Thompson, E. A., Maller, J. L. & Sluder, G. (1999). Requirement of Cdk2-cyclin E activity for repeated centrosome reproduction in Xenopus egg extracts. Science, 283, 851–4.

Holland, A. J., Lan, W., Niessen, S., Hoover, H. & Cleveland, D. W. (2010). Polo-like kinase 4 kinase activity limits centrosome overduplication by autoregulating its own stability. J Cell Biol, 188, 191–8.

Hu, E. & Rubin, C. S. (1990). Casein kinase II from *Caenorhabditis elegans*. Properties and developmental regulation of the enzyme; cloning and sequence analyses of cDNA and the gene for the catalytic subunit. J Biol Chem, 265, 5072–80.

Hu, E. & Rubin, C. S. (1991). Casein kinase II from *Caenorhabditis elegans*. Cloning, characterization, and developmental regulation of the gene encoding the beta subunit. J Biol Chem, 266, 19796–802.

Hu, J., Bae, Y. K., Knobel, K. M. & Barr, M. M. (2006). Casein kinase II and calcineurin modulate TRPP function and ciliary localization. Mol Biol Cell, 17, 2200–11.

Jantsch-Plunger, V., Gonczy, P., Romano, A., Schnabel, H., Hamill, D., Schnabel, R., Hyman, A. A. & Glotzer, M. (2000). CYK-4: A Rho family gtpase activating protein (GAP) required for central spindle formation and cytokinesis. J Cell Biol, 149, 1391–404.

Kachur, T. M., Audhya, A. & Pilgrim, D. B. (2008). UNC-45 is required for NMY-2 contractile function in early embryonic polarity establishment and germline cellularization in *C. elegans*. Dev Biol, 314, 287–99.

Kaitna, S., Mendoza, M., Jantsch-Plunger, V. & Glotzer, M. (2000). Incenp and an aurora-like kinase form a complex essential for chromosome segregation and efficient completion of cytokinesis. Curr Biol, 10, 1172–81.

Kamath, R. S., Fraser, A. G., Dong, Y., Poulin, G., Durbin, R., Gotta, M., Kanapin, A., Le Bot, N., Moreno, S., Sohrmann, M., et al. (2003). Systematic functional analysis of the *Caenorhabditis elegans* genome using RNAi. Nature, 421, 231–7.

Kemp, C. A., Kopish, K. R., Zipperlen, P., Ahringer, J. & O’connell, K. F. (2004). Centrosome maturation and duplication in *C. elegans* require the coiled-coil protein SPD-2. Dev Cell, 6, 511–23.

Kemp, C. A., Song, M. H., Addepalli, M. K., Hunter, G. & O’connell, K. (2007). Suppressors of *zyg-1* define regulators of centrosome duplication and nuclear association in Caenorhabditis elegans. Genetics, 176, 95–113.

Kemphues, K. J., Kusch, M. & Wolf, N. (1988). Maternal-effect lethal mutations on linkage group II of *Caenorhabditis elegans*. Genetics, 120, 977–86.

Kirkham, M., Muller-Reichert, T., Oegema, K., Grill, S. & Hyman, A. A. (2003). SAS-4 is a *C. elegans* centriolar protein that controls centrosome size. Cell, 112, 575–87.

Kitagawa, D., Fluckiger, I., Polanowska, J., Keller, D., Reboul, J. & Gonczy, P. (2011). PP2A phosphatase acts upon SAS-5 to ensure centriole formation in *C. elegans* embryos. Dev Cell, 20, 550–62.

Kitagawa, M. & Lee, S. H. (2015). The chromosomal passenger complex (CPC) as a key orchestrator of orderly mitotic exit and cytokinesis. Front Cell Dev Biol, 3, 14.

Klebba, J. E., Buster, D. W., Nguyen, A. L., Swatkoski, S., Gucek, M., Rusan, N. M. & Rogers, G. C. (2013). Polo-like kinase 4 autodestructs by generating its Slimb-binding phosphodegron. Curr Biol, 23, 2255–61.

Kleylein-Sohn, J., Westendorf, J., Le Clech, M., Habedanck, R., Stierhof, Y. D. & Nigg, E. A. (2007). Plk4-induced centriole biogenesis in human cells. Dev Cell, 13, 190–202.

Kratz, A. S., Barenz, F., Richter, K. T. & Hoffmann, I. (2015). Plk4-dependent phosphorylation of STIL is required for centriole duplication. Biol Open, 4, 370–7.

Lane, H. A. & Nigg, E. A. (1996). Antibody microinjection reveals an essential role for human polo-like kinase 1 (Plk1) in the functional maturation of mitotic centrosomes. J Cell Biol, 135, 1701–13.

Leidel, S., Delattre, M., Cerutti, L., Baumer, K. & Gonczy, P. (2005). SAS-6 defines a protein family required for centrosome duplication in *C. elegans* and in human cells. Nat Cell Biol, 7, 115–25.

Leidel, S. & Gonczy, P. (2003). SAS-4 is essential for centrosome duplication in *C elegans* and is recruited to daughter centrioles once per cell cycle. Dev Cell, 4, 431–9.

Li, H., Liu, X. S., Yang, X., Wang, Y., Wang, Y., Turner, J. R. & Liu, X. (2010). Phosphorylation of CLIP-170 by Plk1 and CK2 promotes timely formation of kinetochore-microtubule attachments. EMBO J, 29, 2953–65.

Lim, A. C., Tiu, S. Y., Li, Q. & Qi, R. Z. (2004). Direct regulation of microtubule dynamics by protein kinase CK2. J Biol Chem, 279, 4433–9.

Meggio, F. & Pinna, L. A. (2003). One-thousand-and-one substrates of protein kinase CK2? FASEB J, 17, 349–68.

Niefind, K., Raaf, J. & Issinger, O. G. (2009). Protein kinase CK2 in health and disease: Protein kinase CK2: from structures to insights. Cell Mol Life Sci, 66, 1800–16.

O’connell, K. F., Caron, C., Kopish, K. R., Hurd, D. D., Kemphues, K. J., Li, Y. & White, J. G. (2001). The *C. elegans zyg-1* gene encodes a regulator of centrosome duplication with distinct maternal and paternal roles in the embryo. Cell, 105, 547–58.

Pagano, M. A., Bain, J., Kazimierczuk, Z., Sarno, S., Ruzzene, M., Di Maira, G., Elliott, M., Orzeszko, A., Cozza, G., Meggio, F., et al. (2008). The selectivity of inhibitors of protein kinase CK2: an update. Biochem J, 415, 353–65.

Peel, N., Dougherty, M., Goeres, J., Liu, Y. & O’connell, K. F. (2012). The *C. elegans* F-box proteins LIN-23 and SEL-10 antagonize centrosome duplication by regulating ZYG-1 levels. J Cell Sci, 125, 3535–44.

Pelletier, L., O’toole, E., Schwager, A., Hyman, A. A. & Muller-Reichert, T. (2006). Centriole assembly in *Caenorhabditis elegans*. Nature, 444, 619–23.

Pelletier, L., Ozlu, N., Hannak, E., Cowan, C., Habermann, B., Ruer, M., Muller-Reichert, T. & Hyman, A. A. (2004). The *Caenorhabditis elegans* centrosomal protein SPD-2 is required for both pericentriolar material recruitment and centriole duplication. Curr Biol, 14, 863–73.

Peng, Y., Wong, C. C., Nakajima, Y., Tyers, R. G., Sarkeshik, A. S., Yates, J., 3rd, Drubin, D. G. & Barnes, G. (2011). Overlapping kinetochore targets of CK2 and Aurora B kinases in mitotic regulation. Mol Biol Cell, 22, 2680–9.

Penner, C. G., Wang, Z. & Litchfield, D. W. (1997). Expression and localization of epitope-tagged protein kinase CK2. J Cell Biochem, 64, 525–37.

Peters, N., Perez, D. E., Song, M. H., Liu, Y., Muller-Reichert, T., Caron, C., Kemphues, K. J. & O’connell, K. F. (2010). Control of mitotic and meiotic centriole duplication by the Plk4-related kinase ZYG-1. J Cell Sci, 123, 795–805.

Praitis, V., Casey, E., Collar, D. & Austin, J. (2001). Creation of low-copy integrated transgenic lines in *Caenorhabditis elegans*. Genetics, 157, 1217–26.

Raich, W. B., Moran, A. N., Rothman, J. H. & Hardin, J. (1998). Cytokinesis and midzone microtubule organization in *Caenorhabditis elegans* require the kinesin-like protein ZEN-4. Mol Biol Cell, 9, 2037–49.

Salvi, M., Raiborg, C., Hanson, P. I., Campsteijn, C., Stenmark, H. & Pinna, L. A. (2014). CK2 involvement in ESCRT-III complex phosphorylation. Arch Biochem Biophys, 545, 83–91.

Salvi, M., Sarno, S., Cesaro, L., Nakamura, H. & Pinna, L. A. (2009). Extraordinary pleiotropy of protein kinase CK2 revealed by weblogo phosphoproteome analysis. Biochim Biophys Acta, 1793, 847–59.

Sarno, S., Reddy, H., Meggio, F., Ruzzene, M., Davies, S. P., Donella-Deana, A., Shugar, D. & Pinna, L. A. (2001). Selectivity of 4,5,6,7-tetrabromobenzotriazole, an ATP site-directed inhibitor of protein kinase CK2 (’casein kinase-2’). FEBS Lett, 496, 44–8.

Sarov, M., Murray, J. I., Schanze, K., Pozniakovski, A., Niu, W., Angermann, K., Hasse, S., Rupprecht, M., Vinis, E., Tinney, M., et al. (2012). A genome-scale resource for *in vivo* tag-based protein function exploration in *C. elegans*. Cell, 150, 855–66.

Schumacher, J. M., Golden, A. & Donovan, P. J. (1998). AIR-2: An Aurora/Ipl1-related protein kinase associated with chromosomes and midbody microtubules is required for polar body extrusion and cytokinesis in *Caenorhabditis elegans* embryos. J Cell Biol, 143, 1635–46.

Severson, A. F., Hamill, D. R., Carter, J. C., Schumacher, J. & Bowerman, B. (2000). The aurora-related kinase AIR-2 recruits ZEN-4/CeMKLP1 to the mitotic spindle at metaphase and is required for cytokinesis. Curr Biol, 10, 1162–71.

Shimada, M., Yamamoto, A., Murakami-Tonami, Y., Nakanishi, M., Yoshida, T., Aiba, H. & Murakami, H. (2009). Casein kinase II is required for the spindle assembly checkpoint by regulating Mad2p in fission yeast. Biochem Biophys Res Commun, 388, 529–32.

Shimanovskaya, E., Viscardi, V., Lesigang, J., Lettman, M. M., Qiao, R., Svergun, D. I., Round, A., Oegema, K. & Dong, G. (2014). Structure of the C. elegans ZYG-1 cryptic polo box suggests a conserved mechanism for centriolar docking of Plk4 kinases. Structure, 22, 1090–104.

Skop, A. R., Liu, H., Yates, J., 3rd, Meyer, B. J. & Heald, R. (2004). Dissection of the mammalian midbody proteome reveals conserved cytokinesis mechanisms. Science, 305, 61–6.

Song, M. H., Aravind, L., Muller-Reichert, T. & O’connell, K. F. (2008). The conserved protein SZY-20 opposes the Plk4-related kinase ZYG-1 to limit centrosome size. Dev Cell, 15, 901–12.

Song, M. H., Liu, Y., Anderson, D. E., Jahng, W. J. & O’connell, K. F. (2011). Protein phosphatase 2A-SUR-6/B55 regulates centriole duplication in *C. elegans* by controlling the levels of centriole assembly factors. Dev Cell, 20, 563–71.

Sonnichsen, B., Koski, L. B., Walsh, A., Marschall, P., Neumann, B., Brehm, M., Alleaume, A. M., Artelt, J., Bettencourt, P., Cassin, E., et al. (2005). Full-genome RNAi profiling of early embryogenesis in *Caenorhabditis elegans*. Nature, 434, 462–9.

St-Denis, N. A., Derksen, D. R. & Litchfield, D. W. (2009). Evidence for regulation of mitotic progression through temporal phosphorylation and dephosphorylation of CK2alpha. Mol Cell Biol, 29, 2068–81.

Strnad, P., Leidel, S., Vinogradova, T., Euteneuer, U., Khodjakov, A. & Gonczy, P. (2007). Regulated HsSAS-6 levels ensure formation of a single procentriole per centriole during the centrosome duplication cycle. Dev Cell, 13, 203–13.

Stubenvoll, M. D., Medley, J. C., Irwin, M. & Song, M. H. (2016). ATX-2, the *C. elegans* Ortholog of Human Ataxin-2, Regulates Centrosome Size and Microtubule Dynamics. PLoS Genet, 12, e1006370.

Szyszka, R., Grankowski, N., Felczak, K. & Shugar, D. (1995). Halogenated benzimidazoles and benzotriazoles as selective inhibitors of protein kinases CK I and CK II from *Saccharomyces cerevisiae* and other sources. Biochem Biophys Res Commun, 208, 418–24.

Toya, M., Iida, Y. & Sugimoto, A. (2010). Imaging of mitotic spindle dynamics in *Caenorhabditis elegans* embryos. Methods Cell Biol, 97, 359–72.

Trembley, J. H., Chen, Z., Unger, G., Slaton, J., Kren, B. T., Van Waes, C. & Ahmed, K. (2010). Emergence of protein kinase CK2 as a key target in cancer therapy. Biofactors, 36, 187–95.

Wang, X., Gupta, P., Fairbanks, J. & Hansen, D. (2014). Protein kinase CK2 both promotes robust proliferation and inhibits the proliferative fate in the *C. elegans* germ line. Dev Biol, 392, 26–41.

Yde, C. W., Olsen, B. B., Meek, D., Watanabe, N. & Guerra, B. (2008). The regulatory beta-subunit of protein kinase CK2 regulates cell-cycle progression at the onset of mitosis. Oncogene, 27, 4986–97.

